# Lysosomal adaptation to extracellular acidity compromises CD8+ T cell cytotoxicity

**DOI:** 10.64898/2026.10.21.760659

**Authors:** Benjamin Segain, Ryan Conesa, Christina Rollings, Sofia Titah, Elise Paulin, Adeline Cozzani, Angelina Kasprowicz, Leila Mohammadnizad, Silvia Gaggero, Yasmine Touil, Ignacio Moraga, Bruno Quesnel, Rahul Roychoudhuri, Robert Eil, Linda V. Sinclair, Jonathan Martinez, Suman Mitra

## Abstract

Extracellular acidosis is a hallmark of aggressive tumors and suppresses cytotoxic-T lymphocyte (CTL)-mediated killing, but how sustained acid exposure reshapes T cell cytotoxicity is poorly understood. Here we show that chronic acidosis drives lysosomal remodeling in human CD8⁺ T cells, converting the lysosome into a proton sink whose adaptation ultimately impairs killing by depleting the intracellular granzyme B reservoir through constitutive vesicle release. Quantitative proteomics of CD8⁺ T cells cultured at pH 6.5 identified lysosomes as the dominant target of acid-driven remodeling: CTLs upregulated the V1 sector of the vacuolar H⁺-ATPase and hyper-acidified the lysosomal lumen. The remodeled compartment accumulated lysosomal membrane proteins (LAMP1, LAMP2, CD63, CD68) post-transcriptionally, acquired higher cathepsin abundance and increased proteolytic activity. Chronic acidity also enlarged endoplasmic reticulum (ER) and lysosomal Ca^2+^ stores without altering cytosolic Ca^2+^, and drives constitutive, Ca^2+^-dependent release of LAMP1^+^ lysosome-derived vesicles, which drains the intracellular reservoir of granzyme B, contributing to reduced CTL activity. These findings establish lysosomal adaptation as a cost of surviving extracellular acidosis and identify lysosomal hydrolase activity as a possible target for improving CTL function in the acidic tumor environment.

## Introduction

The tumor microenvironment (TME) imposes a stable extracellular acidosis on infiltrating immune cells, with interstitial pH typically falling to 6.5 or bellow (Cappellesso et al., 2024; Christiansen et al., 2026; Fischbeck et al., 2020; Gillies et al., 2004). Lactic acid secretion from glycolytic tumor cells and metabolically similar stromal and myeloid populations, primarily drives this acidification (Cappellesso et al., 2024; Vander Heiden et al., 2009). Unlike transient acid shifts at inflammatory sites, tumor acidosis is spatially heterogeneous, poorly buffered, and sustained over days to weeks, so infiltrating lymphocytes encounter it throughout their residence in the tumor (Brand et al., 2016; Christiansen et al., 2026). Notably, acidity acts on T cells independently of direct cell-cell contact, distinguishing it from checkpoint ligand or cell contact-mediated suppression (Fischbeck et al., 2020). Low pH arrests cytotoxic T-lymphocytes (CTL) killing capacity (Calcinotto et al., 2012; Fischbeck et al., 2020), suppresses T cell receptor (TCR)-triggered MAPK signaling and cytokine production (Mendler et al., 2012), and inactivates perforin at the effector step (Hodel et al., 2025). Sustained extracellular acidification also reduces IL-2 responsiveness in CD8^+^ T cells (Gaggero et al., 2022).

Acid-imposed T cell dysfunction is at least partly reversible. Genetic or pharmacological reduction of tumor lactate production restores T and NK cell surveillance in preclinical models (Brand et al., 2016; Navarro et al., 2022; Pilon-Thomas et al., 2016), and systemic buffering with oral bicarbonate improves anti-tumor immune responses and synergizes with checkpoint blockade and adoptive T-cell transfer in mice (Pilon-Thomas et al., 2016). Many in vitro studies have used lactic acid supplementation, making it difficult to separate the effects of low pH from those of lactate as a signaling metabolite. Where this has been tested directly, pH neutralization alone restores CTL function despite continuous lactate exposure (Mendler et al., 2012), indicating that proton concentration is the dominant variable. In the clinical setting, CAR-T cells face the same acidic milieu, where even the manufacturing pH influences their potency and phenotype (Prochazkova et al., 2024) and acid-responsive genetic circuits have been engineered to control CAR-T activity locally (Feng et al., 2026). Acidity is not confined to tumors. Lymph node paracortical zones, where naive T cells are primed by dendritic cells, are profoundly acidic, and T cells themselves generate these niches to constrain their own effector output (Wu et al., 2020). Upon exit to the periphery T cells resume effector capacity, yet re-encounter suppressive acidity upon infiltrating the tumor. Together, these observations establish extracellular acidosis as a pervasive constraint on T cell immunity, yet most functional studies have examined T cell responses over hours of co-culture or acute restimulation. Under prolonged exposure, CD8^+^ T cells adopt a less differentiated, naive-like state with reduced capacity to produce effector cytokines and cytotoxic molecules (Cheng et al., 2023), but how chronic acidosis reshapes the broader molecular landscape of the CD8^+^ T cell has not been systematically examined.

Cells confronting sustained extracellular acidosis must maintain cytoplasmic pH to preserve enzymatic function and survival. Plasma membrane acid extruders, Na⁺/H⁺ exchangers, Na⁺-HCO₃⁻ co-transporters, and monocarboxylate transporters constitute the first line of defense, but intracellular organelles can also serve as proton sinks: the vacuolar H⁺-ATPase (V-ATPase) actively pumps protons into the lysosomal lumen, and increased V-ATPase activity under acid stress could buffer cytoplasmic pH at the cost of further acidifying an already acidic compartment (Liu et al., 2018). V-ATPase activity at the lysosomal membrane also couples proton pumping to metabolic signaling, luminal amino acids signal through the V-ATPase to the Ragulator–Rag complex, recruiting mTORC1 to the lysosomal surface for activation through an inside-out mechanism (Zoncu et al., 2011). Importantly, in tumor cells chronic acid adaptation drives coordinated lysosomal changes and their peripheral redistribution, cathepsin activation, and enhanced exocytic release supporting survival and invasion (Glunde et al., 2003; Robey et al., 2009). In CTLs, the lysosome carries additional significance because lytic granules are secretory lysosomes that share membrane identity, luminal pH, and Ca^2+^-dependent fusion machinery with conventional lysosomes (Blott and Griffiths, 2002; Stinchcombe and Griffiths, 2007); any lysosomal adaptation therefore has direct implications for the killing apparatus. Upon target recognition, TCR triggering mobilizes Ca^2+^ and polarizes these granules toward the immunological synapse, where they undergo Ca^2+^-dependent fusion with the plasma membrane, releasing perforin and granzymes and transiently exposing the granule membrane protein LAMP1 (CD107a) at the cell surface (Stinchcombe and Griffiths, 2007). How extracellular acidity regulates the CTL program remains poorly understood.

Here, we used unbiased proteomics to examine human CD8⁺ T cells adapted to acidic pH and identified the lysosomal compartment as a major target of acid-driven remodeling. Under acidic conditions, lysosomes hyperacidify and trigger Ca^2+^-dependent release of lysosome-derived vesicles, depleting lytic granule cargo before target encounter and dissociating degranulation from effective killing. With prolonged acid exposure, lysosomal membrane proteins accumulate and lysosomal hydrolase activity rises, establishing a broader program of organellar adaptation to chronic extracellular acidosis.

## Results

### Chronic acidity depletes cytotoxic cargo while sparing degranulation

Cytotoxic T cells infiltrating solid tumors are exposed to extracellular pH values around 6.5, yet how chronic acidosis reshapes their cytotoxic program is poorly understood. To address this, we established a chronic acidification model in which anti-CD19 CAR-T cells were cultured in RPMI buffered with PIPES at a stable pH of 7.5 or 6.5 (**Figure 1A and Figure 1 – Figure supplement 1A**). This buffering was required because bicarbonate-buffered medium re-equilibrates with atmospheric CO2 and returns toward neutrality over a multi-day culture, allowing only transient rather than chronic acidification (**Figure 1 – Figure supplement 1B**). CAR-T cells remained viable at pH 6.5 throughout the culture period, with viability comparable to that measured at pH 7.5 (**Figure 1 – Figure supplement 1C**). Although, CAR-T cells expanded less at pH 6.5 than at pH 7.5 (**Figure 1 – Figure supplement 1D**); all subsequent co-cultures were therefore performed at normalized effector-to-target ratios. Surface expression of the anti-CD19 CAR was unchanged between the two conditions (**Figure 1 – Figure supplement 1E**).

**Figure 1.**
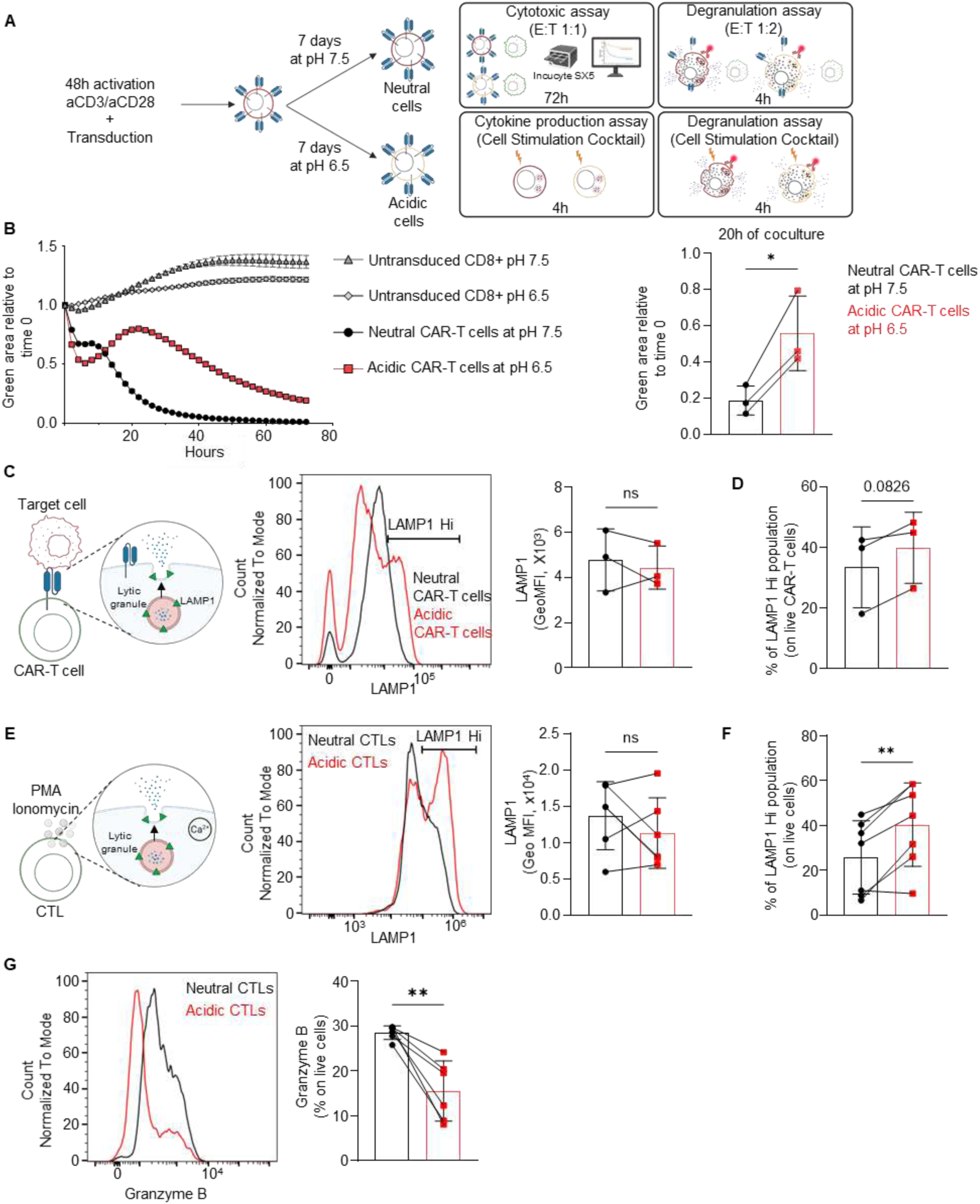
Acidic microenvironment impairs killing activity and reprograms degranulation in human CTLs and CAR-T cells. **(A)** Schematic representation of experimental design, CAR-T cells or CTLs were incubated for 7 days at pH 7.5 or pH 6.5 (neutral and acidic cells) supplemented with IL-2 before functional assays. **(B)** Cytotoxic assay with neutral or acidic CAR-T cells co-cultured with NALM6-GFP (effector:target ratio 1:1) at pH 7.5 or 6.5 for 3 days. GFP signal was measured by IncuCyte SX5. Data are presented as mean +/− SD n= 3 independent donors. **(C left)** Schematic representation of degranulation assay of CAR-T cells co-cultured with target cells. **(C right)** Degranulation assay of neutral or acidic CAR-T cells co-cultured with NALM6-GFP (effector:target ratio 1:2) for 4 h in the presence of anti-human LAMP1 antibody at pH 7.5 or pH 6.5. Data are presented as mean +/− SD n= 3 independent donors. **(D)** Bar graph of LAMP1 population during CAR-T cells degranulation. Data are presented as mean +/− SD n= 3 independent donors. **(E left)** Schematic representation of degranulation assay of CTLs degranulation. **(E right)** Degranulation assay of neutral and acidic CTLs stimulated with Cell Stimulation Cocktail with protein transport inhibitors in different pH for 4h in different pH, and human anti-LAMP1 antibody. Data are presented as mean +/− SD n= 5 independent donors. **(F)** Bar graph of LAMP1 population during CTLs degranulation. Data are presented as mean +/− SD n= 7 independent donors. **(G)** Granzyme B intracellular staining of neutral and acidic CTLs. Data are presented as mean +/− SD n= 6 independent donors. Statistical significance was determined using a paired two-tailed t-test (* p < 0.05, ** p < 0.01, *** p < 0.001). Normal distribution was confirmed by using the Shapiro-Wilk test.

**Figure 1 – Figure supplement 1.**
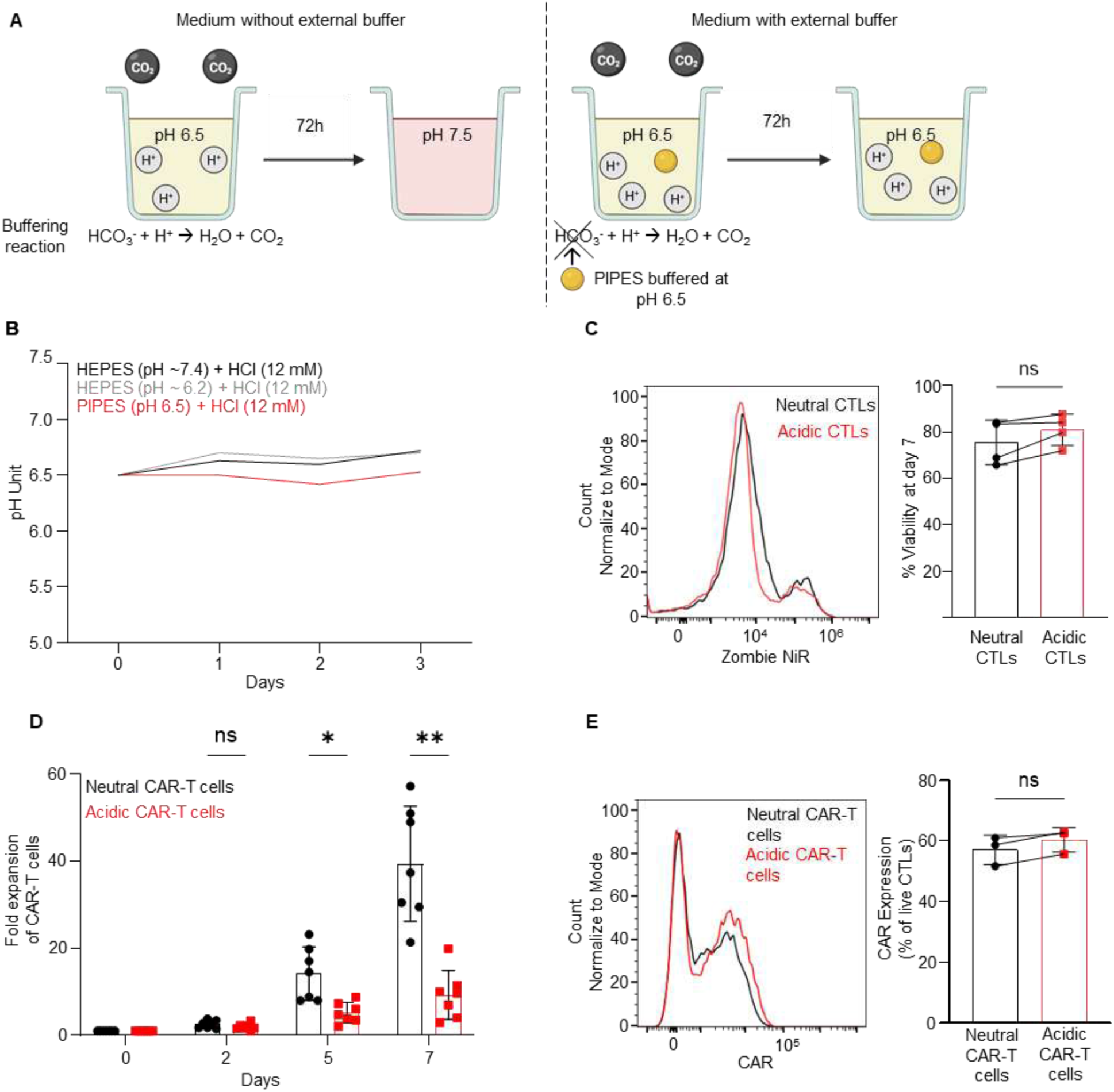
Stable acidic media (pH 6.5) impairs CTLs proliferation without affecting viability and CAR expression. **(A)** Schematic representation of stable acidic media. RPMI media is buffered by sodium bicarbonate to maintain a physiological pH (around 7.2). Adding HCl and PIPES buffer (stabilized at pH 6.5) neutralizes the bicarbonate buffering system, maintaining a stable acidic pH of 6.5. **(B)** pH variation during three days using three different Good’s buffers: HEPES (pH ∼7.4) + 12 mM HCl, HEPES stabilized at pH 6.2 + 12 mM HCl, PIPES stabilized at pH 6.5 + 12 mM HCl. **(C)** Viability of neutral and acidic CTLs after 7 days of culture in respectively pH 7.5 or pH 6.5 (supplemented with IL-2, 20 ng/ml). Data are presented as mean +/− SD, n= 4 independent donors **(D)** Expansion of neutral and acidic CTLs during a week of incubation at pH 7.5 and 6.5 respectively. Data are presented as mean +/− SD, n= 6 independent donors. **(E)** CAR expression in neutral or acidic CTLs after 7 days of culture. Data are presented as mean +/− SD, n= 3 independent donors. Statistical significances were determined using a paired two-tailed t-test (* p < 0.05, ** p < 0.01, *** p < 0.001). Normal distribution was confirmed by using the Shapiro-Wilk test.

**Figure 1 – Figure supplement 2.**
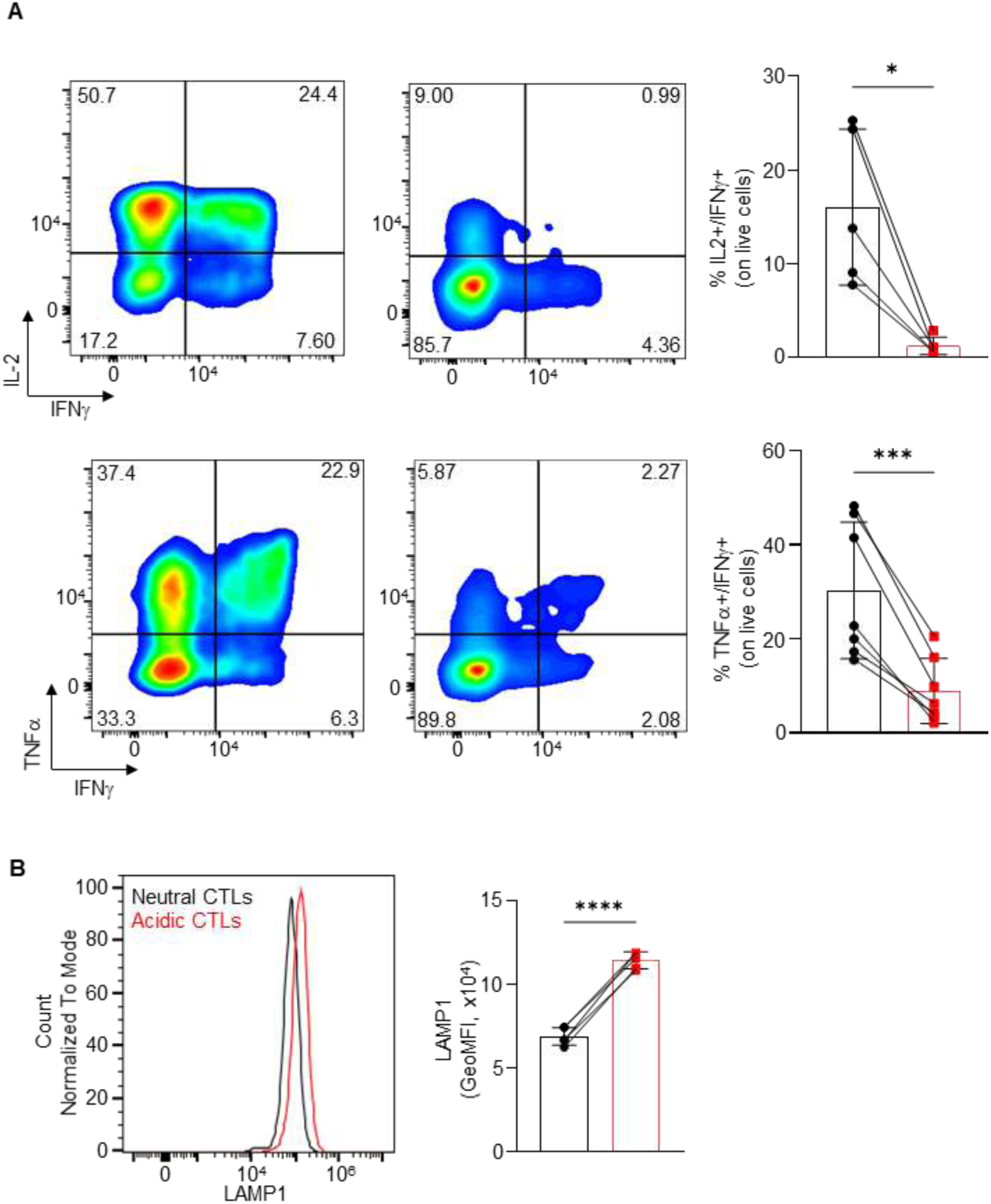
Chronic acidic conditions impair effector function of acidic CTLs. **(A)** Cytokine production assay of neutral or acidic CTL cells stimulated with Cell Stimulation Cocktail plus Golgi inhibitors (eBioscience, 1/500) for 4 h at pH 7.5 or pH 6.5. Data are presented as mean +/− SD, n= 5-7 independent donors. **(B)** Intracellular pool of CD107a in neutral and acidic CTLs. Data are presented as mean +/− SD, n= 5 independent donors. Statistical significances were determined using a paired two-tailed t-test and one way ANOVA test with Tukey’s correction for multiple comparison (* p < 0.05, **** p < 0.001). Normal distribution was confirmed by using the Shapiro-Wilk test.

We then cultured CAR-T cells for 7 days at either pH and measured their cytotoxicity against NALM6 cells in two conditions: cells cultured and assayed at pH 7.5 (neutral CAR-T cells), and cells cultured and assayed at pH 6.5 (acidic CAR-T cells) (**Figure 1A**). This design recapitulates the cytotoxic potential of CAR-T cells chronically grown and maintained in acidic conditions. Untransduced T cell counterparts were included throughout as a control for CAR-independent target death and produced minimal NALM6 killing at either pH (**Figure 1B**), confirming that the cytotoxicity measured here is CAR-dependent. Acidic CAR-T cells tested at pH 6.5 killed NALM6 cells less efficiently than neutral CAR-T cells tested at pH 7.5 (**Figure 1B**). We next asked whether the loss of killing reflected a failure to degranulate, and measured surface exposure of lysosomal membrane protein LAMP1 upon restimulation. During degranulation, secretory lysosomes fuse with the plasma membrane to release their content, resulting in LAMP1 cell surface exposure (Blott and Griffiths, 2002; Peters et al., 1991). Beyond serving as a degranulation marker, this transient surface exposure of LAMP1 protects cytotoxic lymphocytes from perforin-mediated self-damage during target-cell killing (Cohnen et al., 2013). Unexpectedly, LAMP1 levels on acidic CAR-T cells were comparable to those on CAR-T cells expanded at pH 7.5 (**Figure 1C**), indicating that the killing defect is not explained by a failure to mobilize granule membranes. Chronic acidic exposure, drove the emergence of a LAMP1-high population (**Figure 1D**), suggesting that chronic acidic conditions give rise to a CAR-T cell subset with distinct degranulation potential. To assess effector capacity independently of target engagement, we restimulated CTLs with PMA and ionomycin for 4 h and measured cytokine production and degranulation potential by flow cytometry. Production of IFN-γ, TNF-α, and IL-2 was reduced in acidic CTLs tested at pH 6.5 compared with neutral CTLs tested at pH 7.5 (**Figure 1 – Figure supplement 2A**). Membrane LAMP1 levels were similar regardless of culture conditions (**Figure 1E**), yet the LAMP1-high population rose in chronically acid-cultured cells **(Figure 1F)**, confirming the previous observation in CAR-T cells. However, the total amount of LAMP1 available was higher in CTLs chronically cultured in acidic conditions (**Figure 1 – Figure supplement 2B**). Granzyme B content was also reduced in acid-cultured cells (**Figure 1G**), suggesting that chronic acidic conditions deplete pre-formed granzyme B stores, which may reflect a broader reduction in cytotoxic granule cargo. Taken together, these data show that prolonged culture at pH 6.5 impaired CAR-T cell killing, cytokine production, and granzyme B content without a measurable reduction in degranulation as assessed by LAMP1 surface exposure. This dissociation between LAMP1 availability and its mobilization to the plasma membrane, together with the emergence of a LAMP1-high population, suggests that acidity alters the granules themselves rather than the machinery that delivers them, prompting us to define the underlying cellular changes in an unbiased manner.

### Chronic Acidity Remodels the Lysosomal Proteome of Human CTLs

Little is known about how CD8⁺ T cells adapt at the proteome level to acidic environments such as tumors and lymph nodes. To address this, we performed quantitative label-free proteomics on primary human CD8⁺ T cells cultured for 7 days at pH 7.5 or pH 6.5. Four donors were profiled by data-independent acquisition mass spectrometry, quantifying 7,490 protein groups with absolute copy numbers estimated using the proteomic ruler (Wiśniewski et al., 2014). Total protein weight per cell was indistinguishable between conditions (63.7 versus 63.1 pg. **Figure 2 – Figure supplement 1A**), indicating that the observed changes were compositional rather than due to global protein loss. The basal response was compact: 373 proteins changed by more than 1.5-fold (180 downregulated and 193 upregulated in acidic conditions), representing 5% of the quantified proteome (**Figure 2A**, and **Table S1**). Principal-component analysis separated pH conditions along the third component after donor identity dominated the first two, identifying chronic pH exposure as the leading treatment-associated effect on the resting proteome (**Figure 2 – Figure supplement 1A**). Similar numbers of up-and downregulated proteins argue against generalized protein degradation under acidic conditions and instead support coordinated proteomic remodeling.

**Figure 2.**
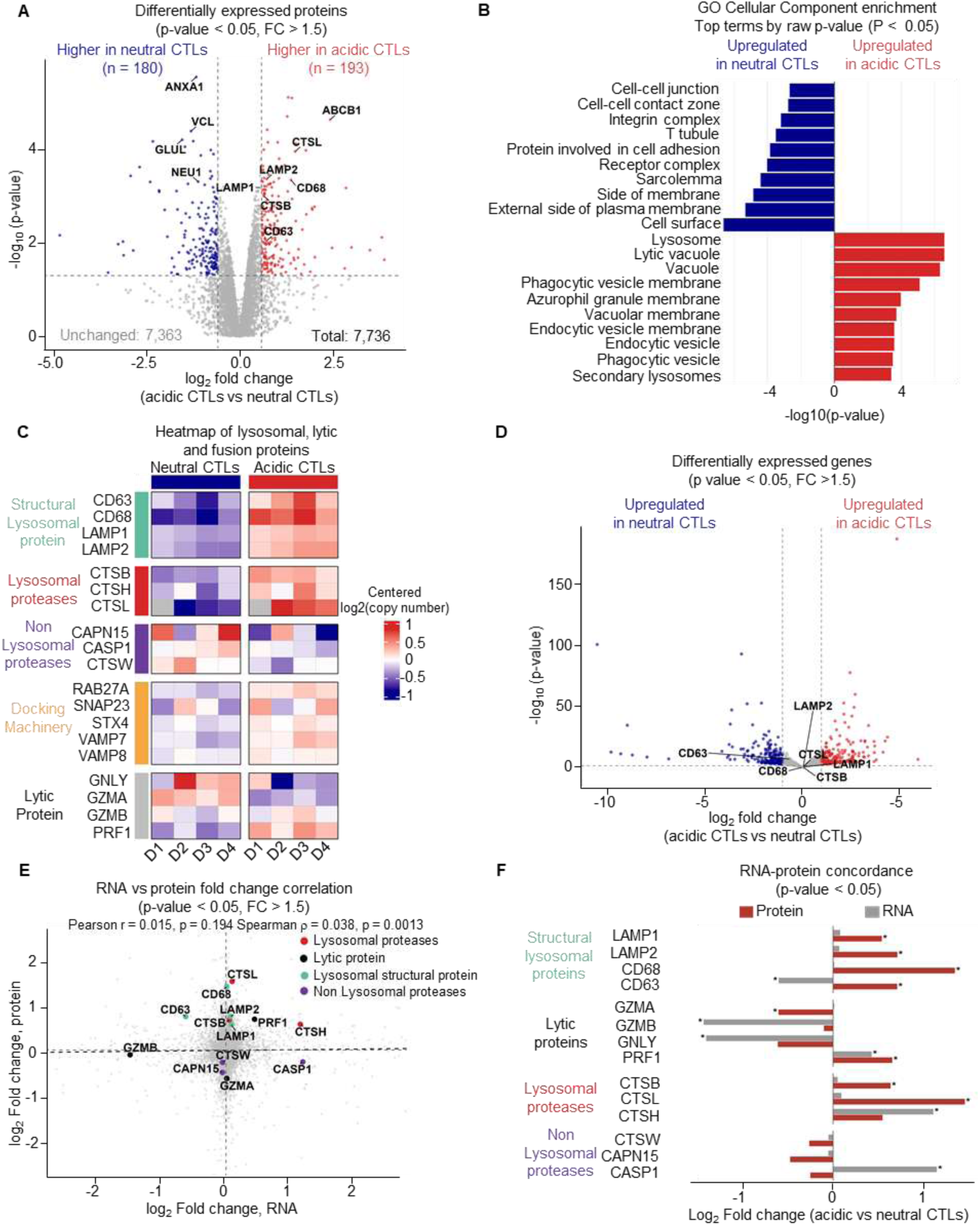
Chronic acidosis remodels CTL lysosomal protein repertoire at a post-transcriptional level. **(A)** Volcano plot of proteomic analysis of acidic CTLs compared to neutral CTLs (7 days at pH 7.5). Dashed lines indicate log₂FC = 0.585 and p < 0.05. **(B)** Top Gene Ontology Cellular Component terms enriched in acidic or neutral CTLs (raw p-value < 0.05). **(C)** Heatmap of structural lysosomal proteins (CD63, CD68, LAMP1, LAMP2), lytic proteins (GNLY, GZMA, GZMB, PRF1), lysosomal proteases (CTSB, CTSH, CTSL), non-lysosomal proteases (CAPN15, CASP1, CTSW) and docking machinery (RAB27A, SNAP23, STX4, VAMP7, VAMP8). Values represent log2-transformed copy numbers centered to each donor’s own mean across conditions. **(D)** Volcano plot of RNA sequencing of acidic CTLs compared to neutral cells. Dashed lines indicate log_2_FC = 0.585 and P < 0.05. **(E)** Scatter plot comparing log_2_ fold change (acidic compared to neutral CTLs) at the RNA level (x-axis) versus the protein level (y-axis). Dashed black line indicates linear regression fit (Pearson r = 0.015, R² < 0.001, p = 0.194); Spearman ρ = 0.038 (p = 0.0013). Structural lysosomal proteins (CD63, CD68, LAMP1, LAMP2), lytic machinery (GZMA, GZMB, PRF1), lysosomal proteases (CTSB, CTSH, CTSL), non-lysosomal hydrolases (CTSW, CAPN15, CASP1) are highlighted. **(F)** Bar plot comparing log_2_ fold change (acidic vs. neutral CTLs) at the protein (red) and RNA (gray) level for structural lysosomal proteins, lytic proteins, lysosomal proteases, and non lysosomal proteases (* p < 0.05). The proteomics data were analyzed using the limpa package. RNA-seq analysis was performed using the DESeq2 package. Proteomics analyses were performed on 4 independent donors.

**Figure 2 – Figure supplement 1.**
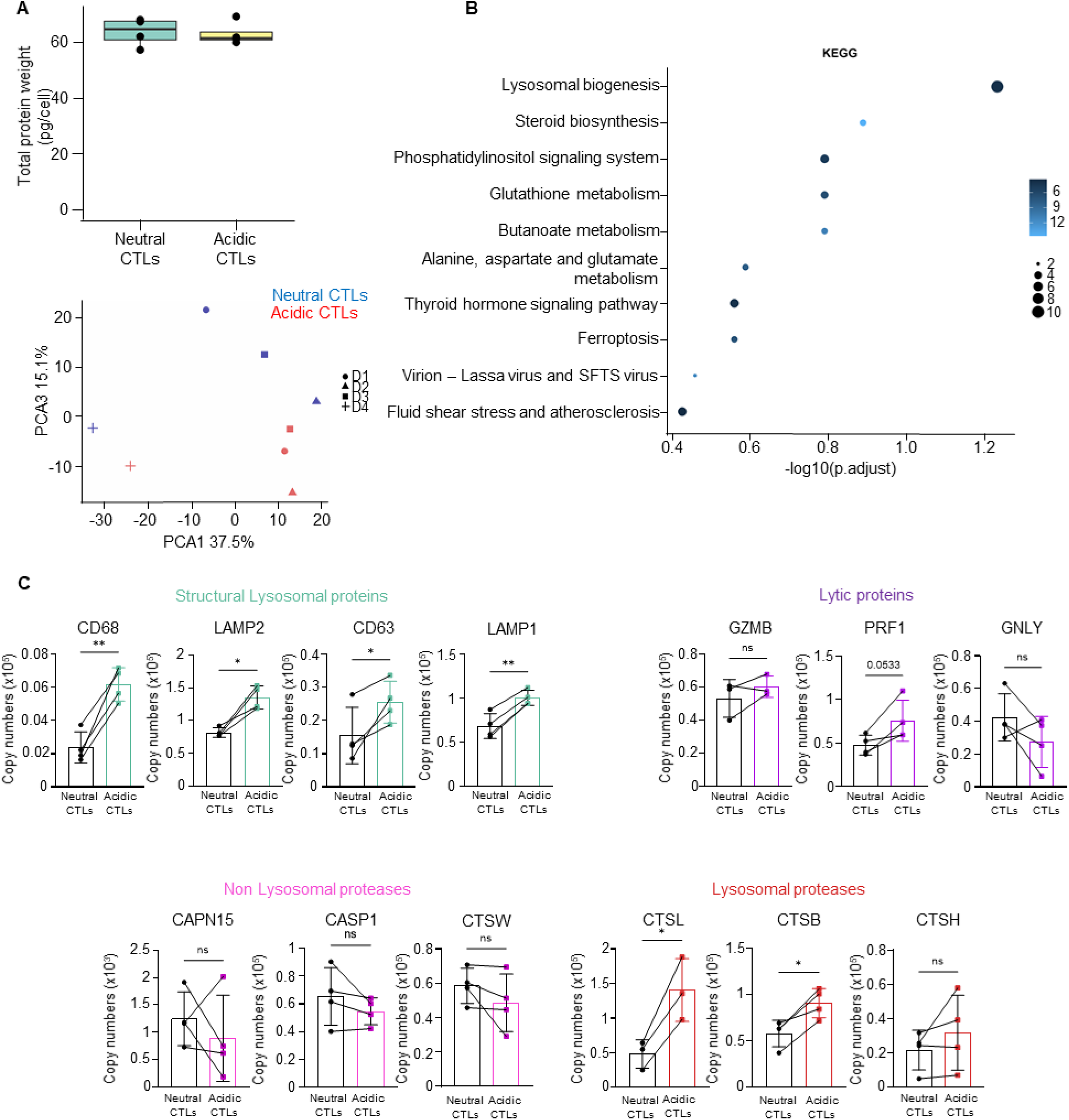
Chronic acidosis switches cathepsin population towards acid-optimized cathepsins. **(A)** Estimated protein weight per cell in neutral and acidic CTLs and principal component analysis with the first component (PCA1, 37.5%) reflecting donor identity and the third component (PCA3, 15.1%) reflecting pH conditions. **(B)** Top 10 Kyoto Encyclopedia of Genes and Genomes (KEGG) terms enriched in acidic CTLs vs neutral CTLs (raw p-value < 0.05). Dot size denotes gene set size and dot color the -log_10_(p-value). **(C)** Individual bar graphs of proteins grouped by their function: structural lysosomal proteins (CD68, LAMP2, CD63, LAMP1), lytic proteins (GZMB, PRF1, GNLY), Pi(3,5)P2 kinase (PIKFYVE), non-lysosomal proteases (CAPM15, CASP1, CTSW), lysosomal proteases (CTSL, CTSB, CTSH). Proteomics analyses were performed on 4 independent donors. Statistical significances were determined using a paired two-tailed t-test (* p < 0.05, ** p < 0.01). Normal distribution was confirmed by using the Shapiro-Wilk test.

**Figure 2 – Figure supplement 2.**
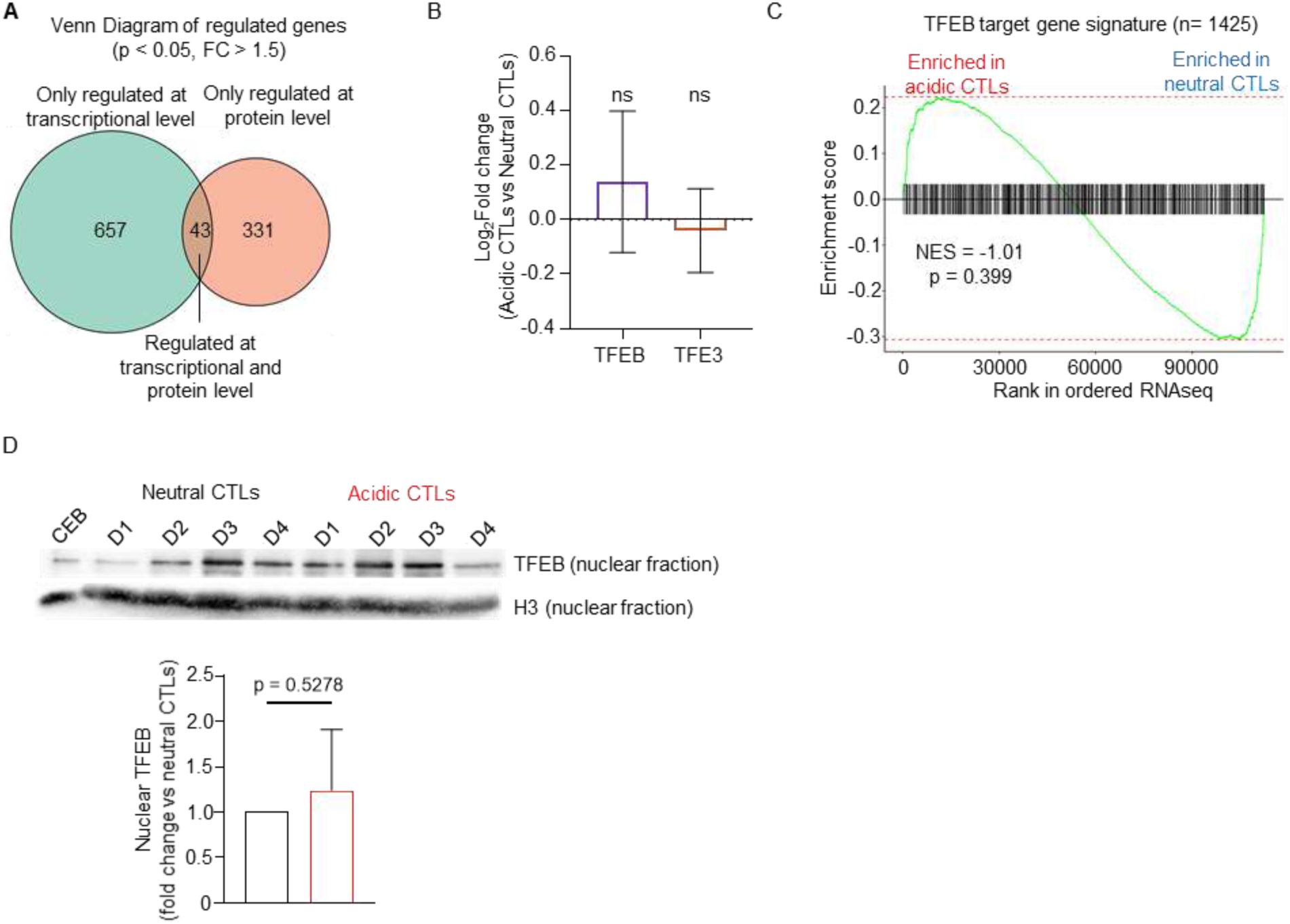
Chronic acidosis remodels the lysosomal protein landscape in a TFEB-independent manner. **(A)** Venn Diagram representing genes significantly regulated (p < 0.05, FC > 1.5) only at RNA level (green, n = 657), at protein level (orange, n = 331) or at both levels (intersection, n = 43). **(B)** TFEB and TFE3 expression in acidic CTLs compared to neutral CTLs. **(C)** Gene Set Enrichment Analysis of TFEB target gene signature (n = 1,425) from MSigDB TFEB_TARGET_GENES gene set (GTRD collection; Yevshin et al., 2018) in acidic CTLs compared to neutral CTLs. **(D)** TFEB nuclear fraction in neutral and acidic CTLs. Data are presented as mean +/− SD, n= 4 independent donors Proteomics analyses were performed on 4 independent donors. Statistical significances were determined using a paired two-tailed t-test (* p < 0.05, ** p < 0.01, *** p < 0.001). Normal distribution was confirmed by using the Shapiro-Wilk test.

Gene Ontology and KEGG pathway enrichment analyses of the changed proteins converged on the lysosomal compartment, which was the top enriched cellular component proteome-wide (**Figure 2B**, **Figure 2 – Figure supplement 1B** and **Table S2, S3**). The enrichment was directional: lysosomal and vacuolar proteins increased while plasma-membrane proteins decreased, suggesting that chronic acidosis redistributes protein mass from the cell surface into the lysosome. Within the compartment, the remodeling was selective. Several lysosomal membrane proteins, including LAMP1, LAMP2, CD68 and CD63, accumulated in acid-cultured CTLs. Interestingly, lysosomal proteases including CTSB, CTSH and CTSL were enriched in acidic CTLs, whereas non-lysosomal proteases such as CAPN15, CASP1 and CTSW were slightly decreased **(Figure 2C**, **Figure 2 – Figure supplement 1C and Table S4)**. This cathepsin switch has been described in tumor cells at low pH (Fehrenbacher and Jäättelä, 2005) but, to our knowledge, not in CTLs. The docking and fusion machinery for lysosomal exocytosis followed the same pattern, with all five quantified effectors increased (RAB27A, VAMP7, VAMP8, SNAP23, STX4). This expanded lysosomal machinery was not matched by its cytotoxic cargo (**Figure 2C and Table S4**).

To ask whether this remodeling was transcriptionally driven, we compared the basal proteome with RNA-sequencing of acidic CTLs and neutral CTLs. The two layers agreed in regulation direction for only 43 of 1,031 regulated proteins **(Figure 2 – Figure supplement 2A, Table S5)**, suggesting that protein-level changes are largely driven by post-transcriptional regulation. Consistently, the lysosomal machinery was largely unchanged at the transcript level: LAMP1, LAMP2, CD63, CD68, CTSB and CTSL despite clear protein accumulation, (**Figure 2D-F and Table S6**). To further confirm our observations, we decided to explore the activation of the MiT/TFE axis, that has been linked to lysosomal biogenesis (Agostini et al., 2022; Settembre et al., 2011; Wang et al., 2020)). A 1,426-gene TFEB-target signature was not induced, and TFEB and TFE3 transcripts themselves were unchanged (**Figure 2 – Figure supplement 2B and 2C** and **Table S7**). Critically, nuclear TFEB were comparable between neutral and acidic CTLs (**Figure 2 – Figure supplement 2D**). Thus, the lysosomal remodeling under chronic acidic conditions reflects post-transcriptional mechanisms rather than a TFEB-driven biogenesis program.

### Chronic Acidity Alters the Early Proteome Response of CTLs to TCR Stimulation

T cell receptor engagement drives a proteome-scale remodeling that builds the metabolic and cytotoxic machinery of the activated T cell. To ask how chronic acid exposure shapes this activation-induced proteome, we performed a second proteomic experiment in an independent cohort of four donors: CD8^+^ T cells cultured for 7 days at pH 7.5 or pH 6.5 were activated for 4 h with anti-CD3/anti-CD28 beads (**Figure 3A and Table S8**). Two pH-switch arms, in which cells were activated at the opposite pH, served to separate the contribution of the adaptation period from the pH present during activation. Between 7,500 and 7,670 proteins were quantified per condition, and protein content was 76 to 80 pg per cell (**Figure 3 – Figure supplement 1A**). Principal-component analysis of the 6,207 proteins quantified in all sixteen samples separated the samples along PC1 (29.9% of variance) by adaptation pH, not by the pH present during activation: cells switched to the opposite pH for the activation window clustered with their adaptation-arm counterparts (**Figure 3B**). The proteome of the activated CTL is therefore dominated by the chronic adaptation state rather than by acute acid exposure. Between the two activated populations, 825 proteins were differentially expressed, of which 610 were more abundant in acid-adapted CTLs (**Figure 3C and Table S9**). An important consideration is whether acid-cultured cells were simply activated less efficiently. The early activation marker CD69 was equally abundant in the two arms (**Figure 3 – Figure supplement 1B**), so the differences described below are not explained by a failure of acid-cultured cells to respond to the stimulus. MYC, which has a very short half-life in T cells and is therefore a sensitive reporter of ongoing signaling (Marchingo et al., 2020; Preston et al., 2015), was also more abundant in acid-cultured cells (**Figure 3 – Figure supplement 1B**). KEGG enrichment analysis of the proteins more abundant at pH 6.5 identified carbon metabolism, branched-chain amino acid degradation, fatty acid degradation, the citrate cycle and the lysosome (**Figure 3D** and **Table S10**). These data show that chronic acidity redirects the metabolic machinery of the activated CTL toward catabolism while the cells remain responsive to antigen receptor engagement.

**Figure 3.**
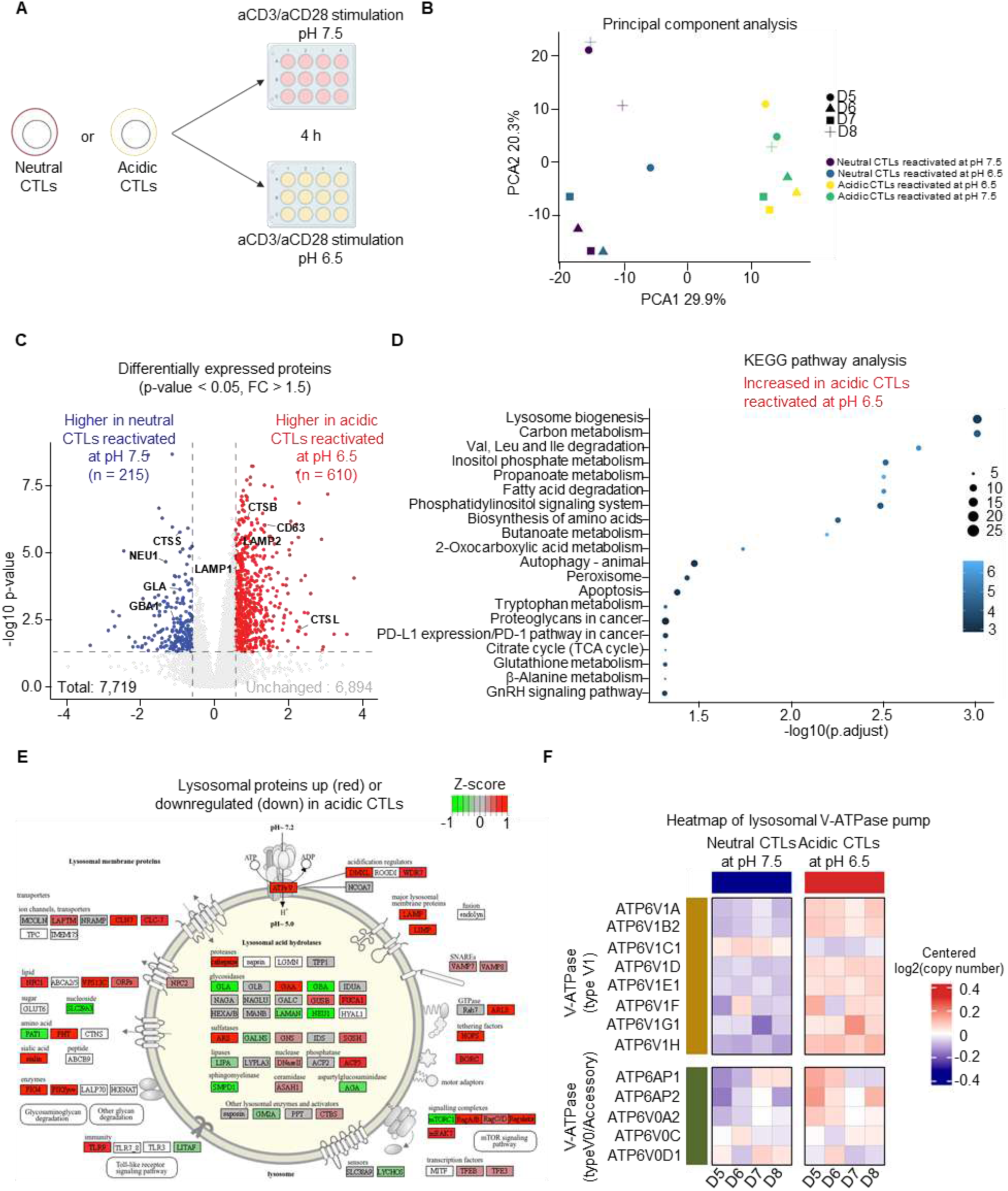
TCR activation of acidic CTL reinforces chronic acidosis lysosomal remodeling. **(A)** Experimental design of TCR activation prior to proteomic analysis of neutral or acidic CTLs**. (B)** Principal component analysis with the first component (PC1, 29.9% of variance) reflecting the adaptation pH and and the second component (PC2, 20.3% of variance) reflecting donor identity. **(C)** Volcano plot of proteomic analysis of acidic CTLs reactivated at pH 6.5 compared to neutral CTLs reactivated at pH 7.5. Dashed lines indicate log₂FC = 0.585 and p < 0.05. **(D)** Top Kyoto Encyclopedia of Genes and Genomes (KEGG) terms enriched in acidic CTLs reactivated at pH 6.5 or neutral CTLs reactivated at pH 7.5 (raw p-value < 0.05). **(E)** KEGG Lysosome biogenesis pathway map at the subcellular level, with proteins colored by log2 fold change of acidic CTLs reactivated at pH 6.5 compared to neutral CTLs reactivated at pH 7.5. Generated using Pathview. **(F)** Heatmap of V-ATPase V1 sector and V-ATPase V0 sector. Values represent log2-transformed copy numbers centered to each donor’s own mean across conditions. Proteomics analyses were performed on 4 independent donors.

**Figure 3 – Figure supplement 1:**
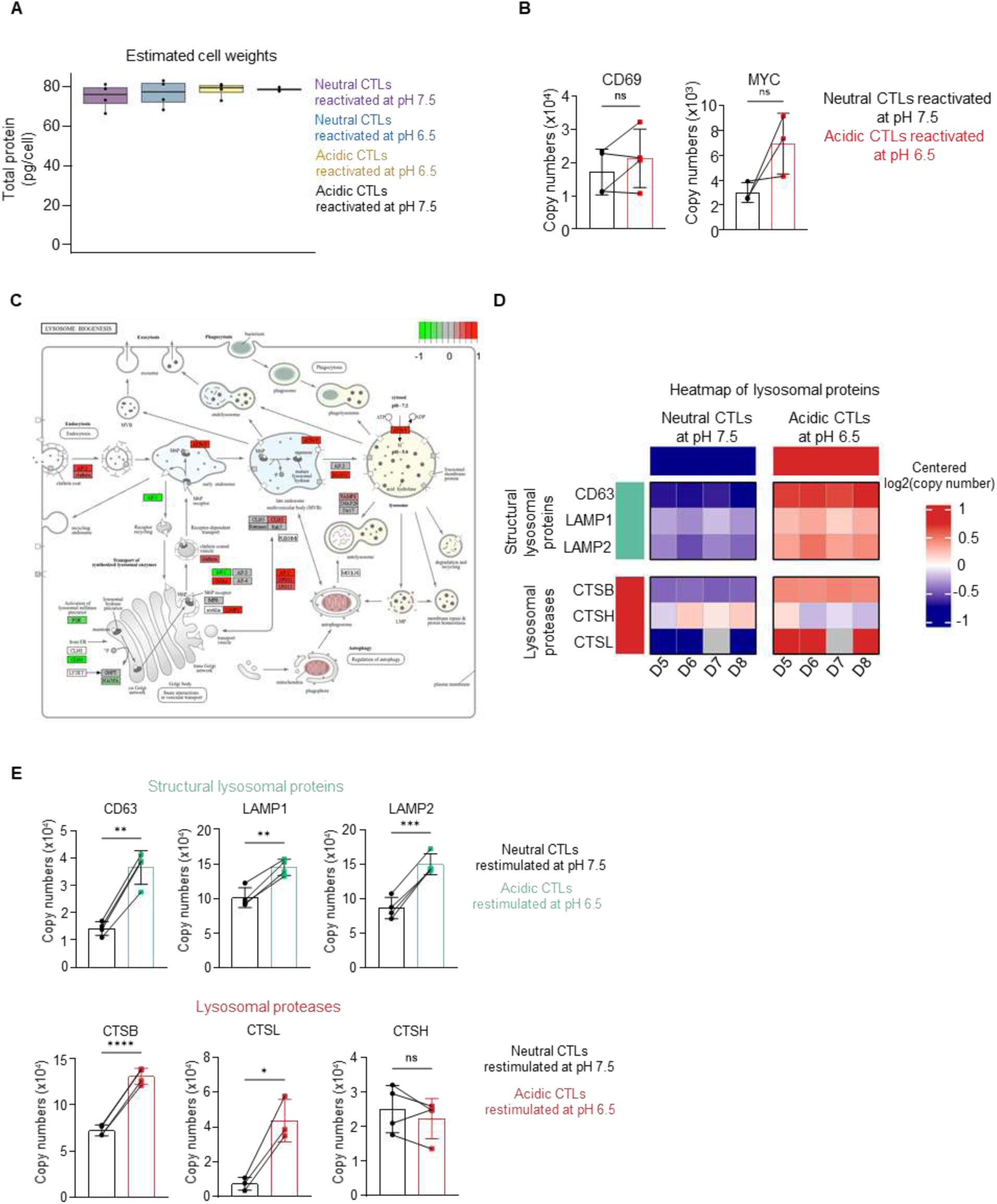
Chronic acidosis reshapes lysosomal biogenesis pathway in TCR-activated CTLs. **(A)** Estimated protein weight per cell in neutral and acidic CTLs reactivated at pH 7.5 or pH 6.5, respectively. **(B)** Bar graphs of CD69 and MYC proteins in neutral CTLs reactivated at pH 7.5 and acidic CTLs reactivated at pH 6.5. **(C)** KEGG lysosome biogenesis pathway at the subcellular level with proteins colored by log2 fold change of acidic CTLs reactivated at pH 6.5 compared to neutral CTLs reactivated at pH 7.5. Generated using Pathview. **(D)** Heatmap of structural lysosomal proteins (CD63, CTSB, CTSL, LAMP1, LAMP2) and lysosomal proteases (CTSS, GBA1, NEU1, GLA). Values represent log2-transformed copy numbers centered to each donor’s own mean across conditions. **(E)** Bar graphs of individual proteins in neutral CTLs reactivated at pH 7.5 and acidic CTLs reactivated at pH 6.5, grouped by function: structural lysosomal proteins (CD63, LAMP1 and LAMP2), lysosomal proteases (CTSL, CTSB, CTSH). Proteomics analyses were performed on 4 independent donors. Statistical significances were determined using a paired two-tailed t-test (* p < 0.05, ** p < 0.01, *** p < 0.001). Normal distribution was confirmed by using the Shapiro-Wilk test.

**Figure 3 – Figure supplement 2:**
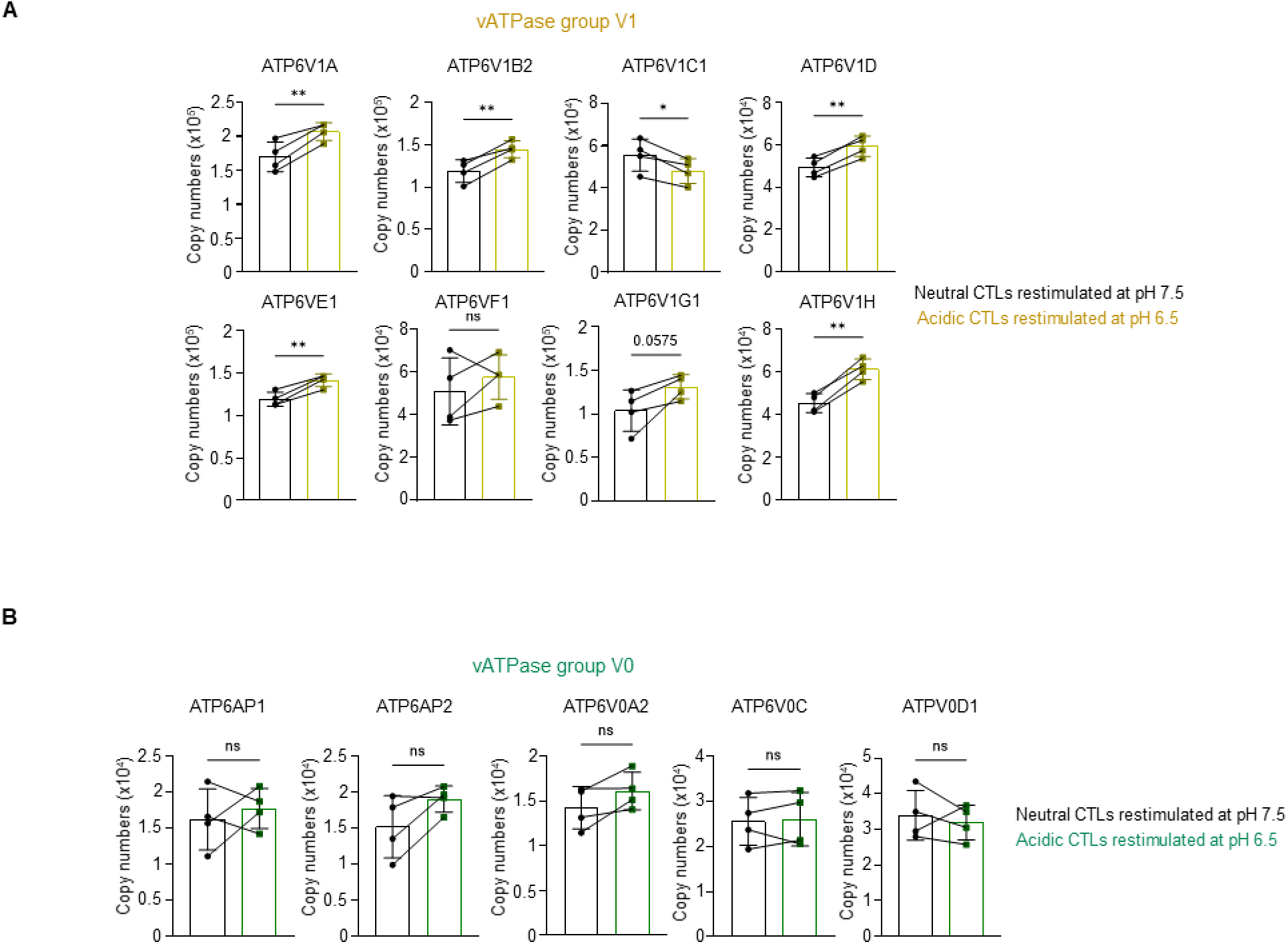
Chronic acidosis reshapes V-ATPase expression in TCR-activated CTLs. **(A)** Bar graphs of individual V-ATPase subunits V1 and **(B)** V0 in neutral CTLs reactivated at pH 7.5 and acidic CTLs reactivated at pH 6.5. Proteomics analyses were performed on 4 independent donors. Statistical significances were determined using a paired two-tailed t-test (* p < 0.05, ** p < 0.01, *** p < 0.001). Normal distribution was confirmed by using the Shapiro-Wilk test.

The lysosomal compartment was enriched among the proteins that both increased and decreased at pH 6.5, reproducing in reactivated cells the directional split seen without restimulation, and the same proteins were involved (**Figure 3E and Figure 3 – Figure supplement 1C-E**). Cathepsin L increased from ∼7,400 to ∼35,700 copies per cell, cathepsin B from ∼72,000 to ∼131,000, CD63 from ∼14,000 to ∼37,000, LAMP2 from ∼87,000 to ∼150,000 and LAMP1 from ∼102,000 to ∼145,000 (**Figure 3E** and **Figure 3 – Figure supplement 1C-E**). Acidic CTLs also contained more of the vacuolar H⁺-ATPase, the proton pump that acidifies the lysosomal lumen. Of 13 subunits quantified, 7 were more abundant in acid-cultured cells, and the increase was restricted to the peripheral V1 sector that hydrolyzes ATP (**Figure 3F** and **Figure 3 – Figure supplement 2A**). ATP6V1A increased from ∼170,000 to ∼207,000 copies per cell and ATP6V1B2 from ∼119,000 to ∼145,000, whereas the membrane-embedded V0 subunits were unchanged (**Figure 3F and Figure 3 – Figure supplement 2B**). Acid-cultured CTLs therefore carry a larger V1 sector available for assembly onto lysosomal V0, which would be predicted to further acidify the lysosomal lumen, that we test directly below.

### Acid-Cultured CTLs Harbor Hyperacidified and Proteolytically Active Lysosomes

To validate the proteomic remodeling by an independent approach, we immunoblotted lysates from CTLs cultured for 3 days at pH 7.5 or pH 6.5. The lysosomal membrane proteins LAMP1 and LAMP2 were increased in acidic CTLs, consistent with the proteomic data (**Figure 4A**). We extended this validation to lysosomal hydrolases and confirmed the accumulation of cathepsins D and L at acidic pH (**Figure 4B**). Unexpectedly, cathepsin B protein levels decreased in acidic conditions, contrasting with the increase observed in the proteomic analysis (**Figure 4B**). This discrepancy may reflect donor-dependent variability or differences in techniques.

**Figure 4.**
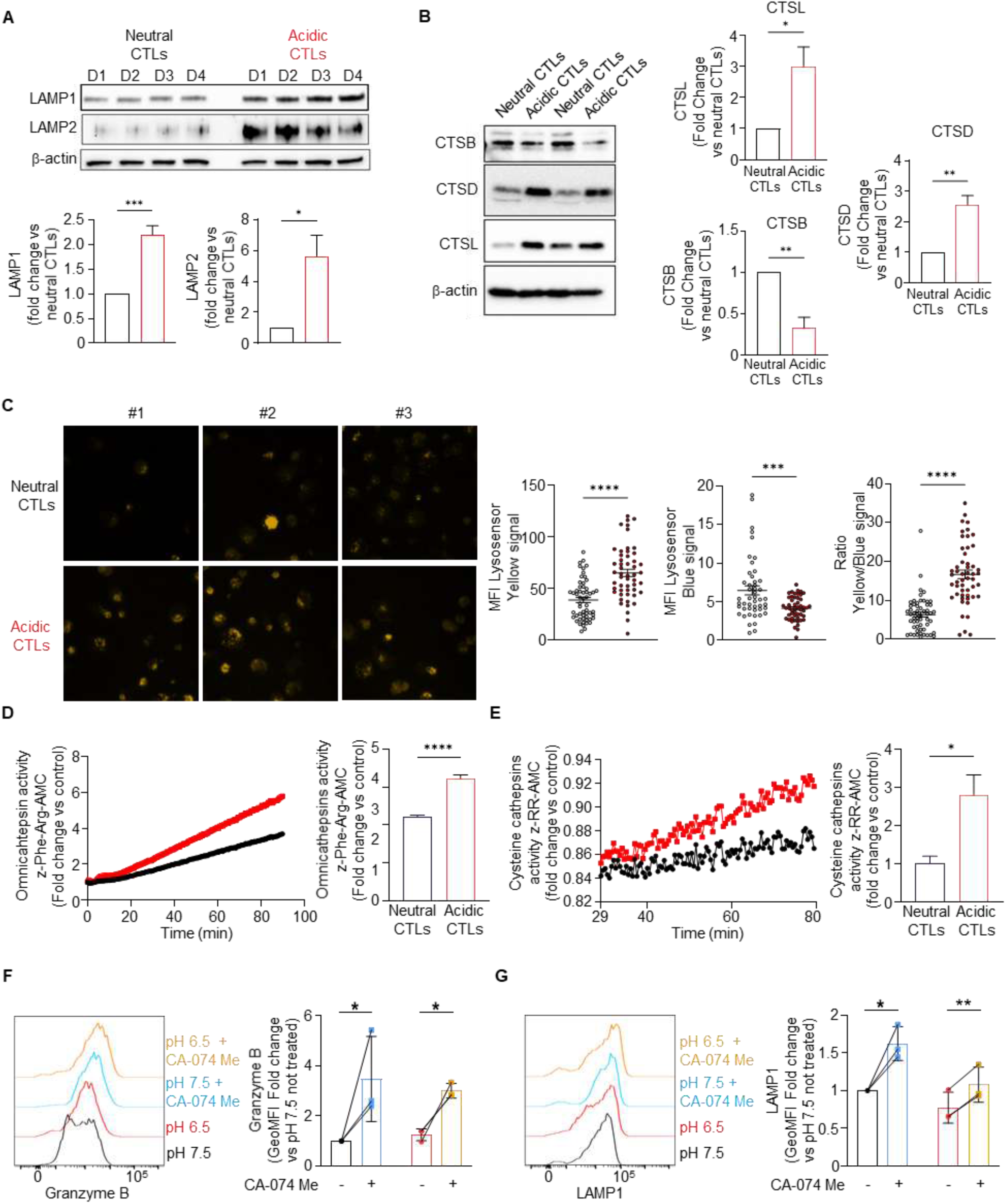
Chronic acidosis promotes hyperacidification and cathepsin hyperactivity in acidic CTLs. **(A)** Levels of LAMP1, LAMP2, **(B)** CTSB, CTSD, CTSL were measured in neutral and acidic CTLs by western blot. β-actin was used as loading control. Data are presented as mean +/− SEM, n= 4 independent donors. **(C)** Ratiometric LysoSensor Yellow/Blue imaging of neutral and acidic CTLs. Data are presented as mean +/− SEM, n= 4 independent donors. **(D)** Omnicathepsins (Z-Phe-Arg-AMC substrate) and **(E)** Cysteine cathepsins (Z-Arg-Arg-AMC substrate) activity was measured in neutral and acidic CTLs. Data are presented as mean only, n= 4 independent donors. **(F)** Granzyme B intracellular staining of CTLs grown 3 days at pH 7.5 or 6.5 in the presence or not of CA074Me (10 µM). Data are presented as mean +/− SD, n= 4 independent donors. **(G)** Degranulation assay of 3 days cultured CTLs in neutral or acidic conditions supplemented by Ca-074 Me stimulated with Cell Stimulation Cocktail in different pH for 4h, in the presence of protein transport inhibitors and human anti-LAMP1. Data are presented as mean +/− SD, n= 4 independent donors. Statistical significance was determined using an unpaired t-test, (* p < 0.05, ** p < 0.01, *** p < 0.001, **** p < 0.0001). For (F), statistical significance was determined using a paired t-test on log2(fold change) (* p < 0.05).

**Figure 4 – Figure supplement 1.**
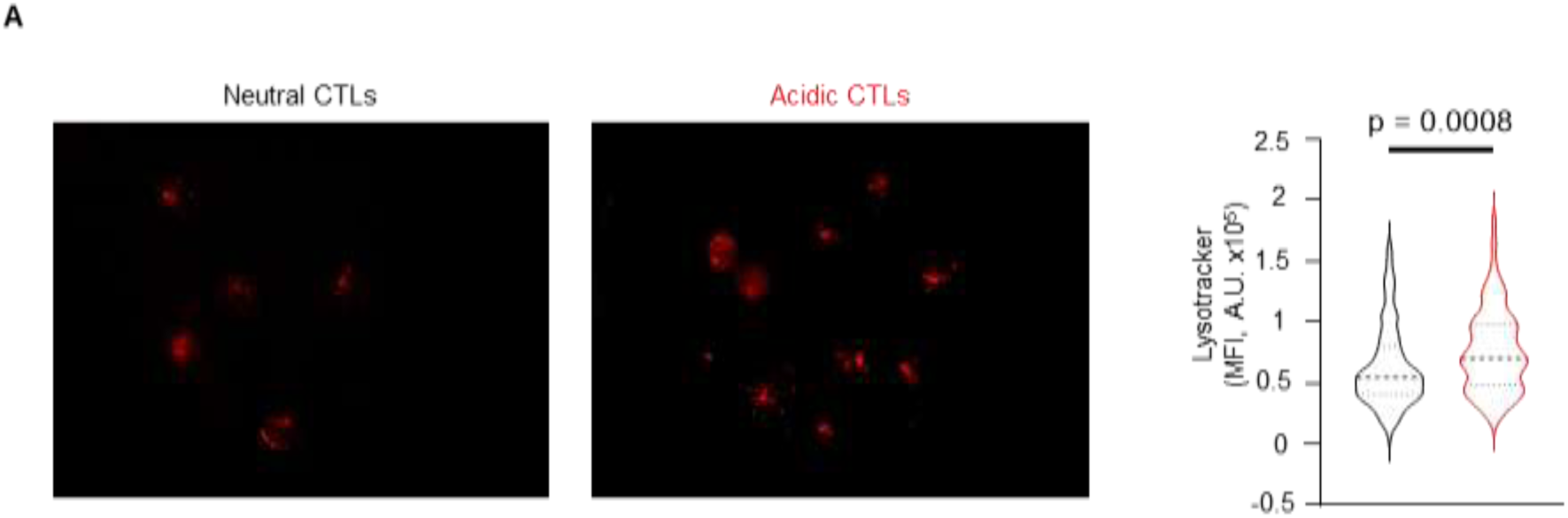
Chronic acidic conditions slightly increase acidic organelles in CTLs. **(A)** LysoTracker signal in neutral and acidic. Lysotracker signal was normalized to cell area and background removal. Data are presented with median (thick dotted line) and first and third quartiles (dashed line), n= 4 independent donors. Statistical significances were determined using Welsh’s t-test.

The coordinated increase in lysosomal protease abundance raised the question of how lysosomes themselves are organized in acidic CTLs. To address this, we used two complementary imaging assays. First, we stained cells with LysoTracker, which labels acidic intracellular organelles. To avoid direct effects of extracellular pH on dye loading, cells from each condition were labeled at neutral pH. LysoTracker signal slightly but significantly increased in acid-cultured cells relative to controls (**Figure 4 – Figure supplement 1A**), indicating that the acidic compartment persists in acidic CTLs. Because LysoTracker accumulation depends jointly on the number, volume and luminal acidity of acidic organelles, this measurement alone cannot distinguish between fewer compartments and a change in their properties. We therefore performed ratiometric LysoSensor Yellow/Blue imaging, which reports luminal pH independently of organelle number. Acid-cultured CTLs displayed a higher Yellow/Blue ratio, indicating a more acidic lysosomal lumen (**Figure 4C**), consistent with the increased abundance of V-ATPase subunits in the proteomic data. Together, these data show that the LysoTracker-positive compartment is maintained in acidic CTLs and that its lumens are hyperacidified, in line with the compositional remodeling of the lysosomal compartment evident in the proteomic data.

To test whether the protein-level changes and lysosome hyperacidification translate into altered proteolytic function, we measured lysosomal protease activity in total lysates. By immunoblotting, we validated the accumulation of CTSL and CTSD (**Figure 4B**), and two fluorogenic substrates further confirmed an increase in omni-cathepsin and cysteine cathepsin activities, indicating a higher proteolytic capacity of the lysosomal compartment in acidic CTLs (**Figure 4D, E**). Cysteine cathepsins are pH-sensitive proteases with an acidic pH optimum and are irreversibly inactivated near neutral pH (Turk et al., 1999, 2012). Because cysteine cathepsins require an acidic lumen, the hyperacidified compartment described above provides a mechanistic basis for this increased proteolytic capacity. We next asked whether this activity contributes to the depletion of lytic cargo in acid-cultured CTLs. CTLs were cultured at pH 7.5 or pH 6.5 for 3 days in the presence of Ca074-Me, a cell-permeant irreversible inhibitor that inactivates cathepsins B and L (Buttle et al., 1992; Montaser et al., 2002). Cathepsin inhibition increased intracellular granzyme B content at both pH values (**Figure 4F**), indicating that cysteine cathepsin activity tonically limits the granzyme B content of CTLs. We next asked whether cathepsin activity also controls the distribution of lysosomal membrane proteins. Following T cell activation with PMA and Ionomycin, surface LAMP1 induction was slightly attenuated in acid-cultured CTLs (**Figure 4G**). Ca074-Me treatment increased activation-induced surface LAMP1 in both neutral and acidic pH (**Figure 4G**), indicating that cysteine cathepsins activity depletes the lysosomal membrane protein pool available for activation-induced surface exposure. Together with the accumulation of LAMP1 in secreted vesicles (**Figure 5A**), these data show that chronic acidity redistributes LAMP1 among three fates, intracellular accumulation, constitutive vesicular release, and attenuated activation-induced surface exposure.

**Figure 5.**
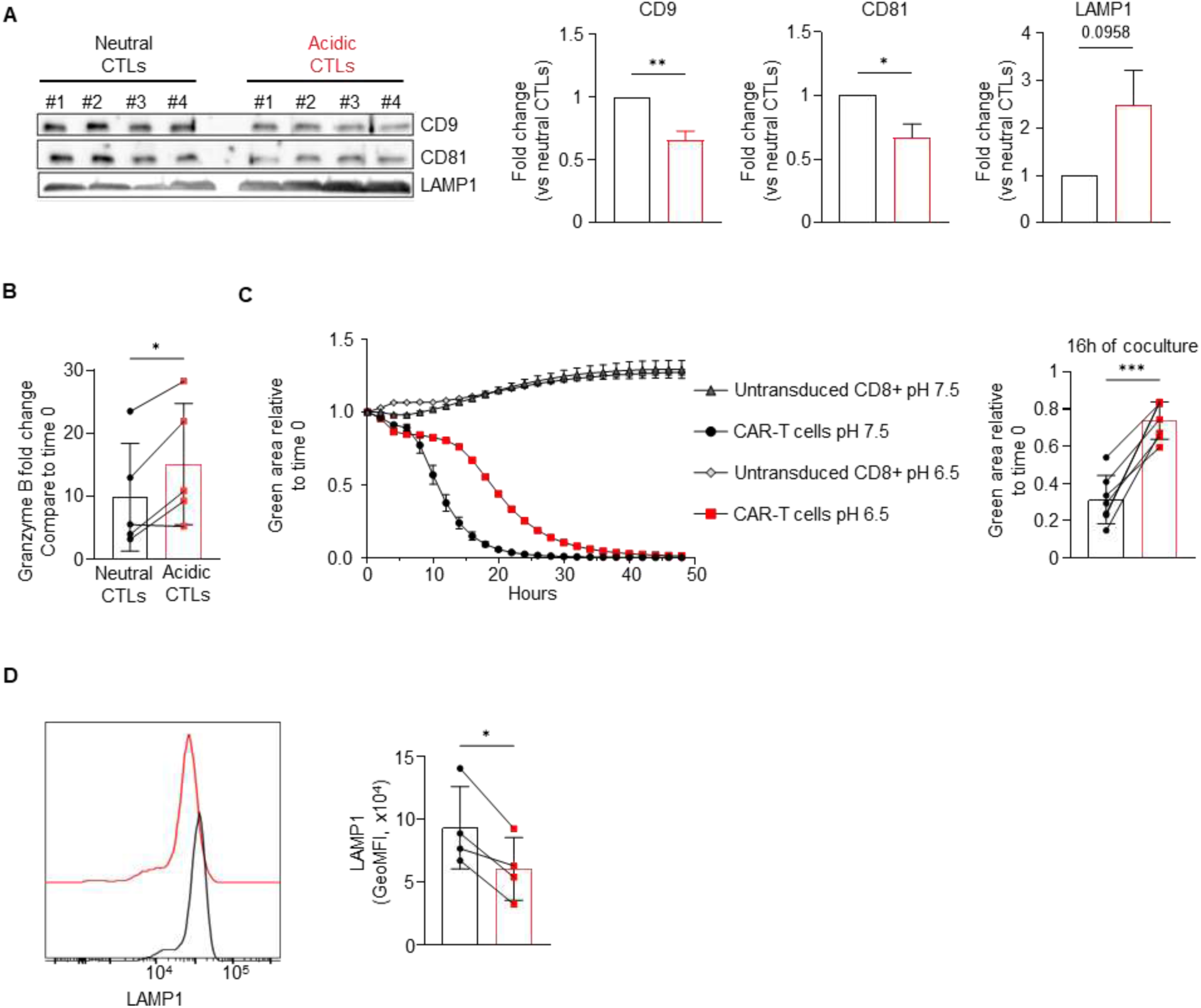
Chronic acidosis triggers constitutive extracellular release of lysosomal cargo. **(A)** Levels of CD9, CD81 and LAMP1 assessed by western blot on culture supernatants of neutral or acidic CTLs three days after aCD3/aCD28 activation. Quantity loaded was normalized based on cell count. Data are presented as mean +/− SEM, n= 4 independent donors. (**B)** Extracellular granzyme B concentration fold change measured by ELISA in the culture supernatant of CTLs stimulated with Cell Stimulation Cocktail for 4 h at pH 7.5, washed and then incubated overnight at pH 7.5 or pH 6.5. Data are presented as mean +/− SD, n= 5 independent donors. **(C)** Cytotoxic assay with CTL CAR-T cells co-cultured with NALM6-GFP (effector:target ratio 1:1) at pH 7.5 or 6.5 for 2 days. GFP signal was measured by IncuCyte SX5. Data are presented as mean +/− SD, n= 7 independent donors. **(D)** Degranulation assay of CTLs stimulated with Cell Stimulation Cocktail for 4h in different pH, in the presence of protein transport inhibitors and human anti-LAMP1. Data are presented as mean +/− SD, n= 4 independent donors. Statistical significance was determined by using an unpaired t-test for western-blot quantification and paired two-tailed t-test for the cytotoxic activity and degranulation assay (* p < 0.05, ** p < 0.01, *** p < 0.001, **** p < 0.0001). Normal distribution was confirmed by using the Shapiro-Wilk test.

### Acidic extracellular pH drives continuous lysosomal exocytosis

Because chronic acidic culture increased intracellular LAMP1 levels without a corresponding increase in LAMP1 at the plasma membrane, we asked whether acidic stress drives constitutive release of lysosomal cargo in the absence of a target. To distinguish a selective cellular response from passive vesicular leakage, we profiled conditioned media from acid-cultured and control CTLs by western blot, targeting markers of distinct secretory pathways: the tetraspanins CD9 and CD81 for multivesicular body (MVB)–derived exosome release (Rydland et al., 2023), LAMP1 for lysosome-derived exosomes release (Shen et al., 2026). Acid-conditioned CTLs showed reduced extracellular CD9 and CD81 compared to controls, ruling out a nonspecific increase in MVB-derived exosome secretion. LAMP1, by contrast, accumulated in acidic supernatants (**Figure 5A**). The release driven by acidity was therefore selective for lysosome-derived vesicles.

Given that acid-conditioned CTLs display a LAMP1-high, granzyme B–low intracellular phenotype and increased lysosome-derived vesicle release, we next asked whether acidic pH forces continuous release of granzyme B after cessation of an activation signal. CTLs were stimulated with PMA and ionomycin for 4 hours at pH 7.5, the stimulus washed out, and the cells incubated overnight at either pH 7.5 or pH 6.5. Extracellular granzyme B was significantly greater at acidic pH (**Figure 5B**), yet intracellular granzyme B was lower in acidic CTLs (**Figure 1G**). Acidic pH thus drives lysosomal exocytosis of lytic cargo that persists after stimulus removal. To test the functional consequence of this constitutive release, we performed a cytotoxic assay in which CAR-T cells that had never experienced acidic conditions were co-cultured with target cells at pH 6.5. CAR-T cell cytotoxic activity decreased in acidic conditions (**Figure 5C**) indicating that even acute acid exposure is sufficient to impair killing. Additionally, acute acidic conditions decrease the degranulation capacity of CTLs (**Figure 5D**). Because this exocytic process relies on intracellular Ca^2+^ mobilization, we next asked whether chronic acidic conditions alter the Ca^2+^ stores that could drive the constitutive release.

### Chronic acidosis enlarges the lysosomal Ca^2+^ store

Lysosomal exocytosis is a Ca^2+^-dependent process (Hessvik and Llorente, 2017; Savina et al., 2003). Because acidic CTLs show increased lysosome-derived exosome secretion, we asked whether chronic acidosis alters the intracellular Ca^2+^ stores that drive this process. CTLs cultured for 7 days at pH 7.5 or pH 6.5 were loaded with the ratiometric Ca^2+^ indicator Indo-1 AM and recorded in Ca^2+^-free medium to isolate intracellular stores from extracellular Ca^2+^ entry. Cells were sequentially challenged with thapsigargin to release the ER pool, GPN to release the lysosomal pool, and ionomycin to mobilize total cellular Ca2+ (**Figure 6A**). Basal cytosolic Ca^2+^ was defined as the mean Indo-1 ratio during the first 120 seconds of recording; organellar Ca^2+^ release was calculated as the difference between the peak ratio after each stimulus and the preceding baseline. Both the ER- and lysosome-releasable stores were significantly enlarged in acidic CTLs (**Figure 6B** and **6C**). The magnitude of store enlargement followed the expected hierarchy of compartmental Ca^2+^ content, with the ER pool exceeding the lysosomal pool in both conditions. Basal cytosolic Ca^2+^ was similar between conditions, and total ionomycin-releasable Ca^2+^ showed only a non-significant trend toward increase (**Figure 6D**). Chronic acidosis thus enlarges the releasable ER and lysosomal Ca²⁺ stores of CTLs, providing a mechanism through which acid conditions could sustain the constitutive lysosome-derived vesicle release observed above.

**Figure 6.**
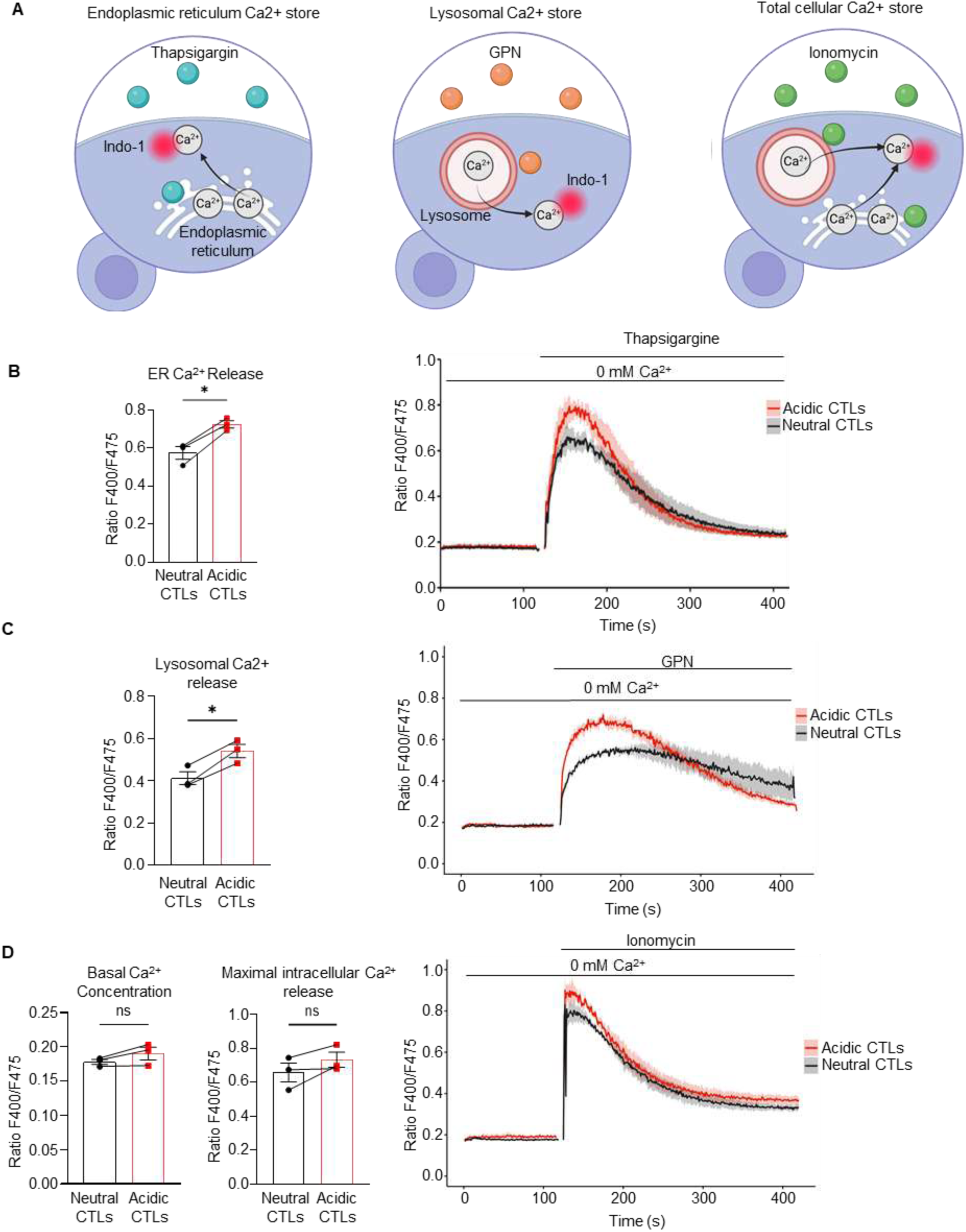
Chronic acidosis enlarges lysosomal Ca²⁺ stores. **(A)** Schematic representation of the different molecules used to explore Ca^2+^ stores. Thapsigargin for endoplasmic reticulum, glycyl-L-phenylalanine 2-naphthylamide (GPN) for Ca^2+^ lysosomal stores, Ionomycin for total Ca^2+^ mobilization. Data are presented as mean +/− SEM, n= 5 independent donors. **(B)** Indo-1 AM calcium flux traces in neutral and acidic CTLs, stimulated in sequence with thapsigargin (1 µM). Data are presented as mean +/− SEM, n= 5 independent donors. **(C)** glycyl-L-phenylalanine 2-naphthylamide (50 µM), and **(D)** ionomycin (10 µM), in 0 mM extracellular Ca²⁺. Data are presented as mean +/− SEM, n= 5 independent donors. Statistical significance was determined by paired two-tailed t-test

Collectively, these findings demonstrate that human CD8⁺ T cells respond to sustained extracellular acidosis through a coordinated lysosomal reprogramming in which the lysosome serves as a proton sink, driving luminal hyper acidification and a shift toward cathepsin activity that degrades cytotoxic cargo; accordingly, pharmacological inhibition of cysteine cathepsins increased intracellular granzyme B content and surface LAMP1 expression in acidic CTLs, establishing lysosomal proteolysis as a targetable vulnerability of the acid-adapted state.

## Discussion

The present study identifies the lysosome as a central hub for the adaptation of human CD8⁺ T cells to chronic extracellular acidosis. The fundamental finding is that sustained exposure to acid pH reprograms the lysosomal compartment of CTLs.

Our results indicate that both acute and chronic acid conditions drastically reduce CTL killing capacity by triggering untargeted release of lysosome-related organelles. Specifically, we demonstrate how the lysosomal compartment under acidic conditions acts as a proton sink through the upregulation of the V-ATPase V1 sector, resulting in hyperacidification of the intraluminal pH of the lysosome and increased proteolytic capacity. Additionally, chronic acidic conditions lead to an increase in the levels of Ca^2+^ within the ER and the lysosomal compartment. Increased luminal Ca^2+^ levels have been shown to increase the sensitivity of Ca^2+^ stores to IP3 (Nunn and Taylor, 1992), promoting exocytosis. Thus, our data show that acid conditions result in an imbalance in the intracellular Ca^2+^ storage, leading to untargeted lysosome-related organelles secretion, reduced lytic granule content in CTLs and therefore explaining the reduced killing capacity of CTLs under acid conditions. Exocytosis is a Ca^2+^-dependent process (Hessvik and Llorente, 2017; Savina et al., 2003), and two-pore channels on cytotoxic granules promote their release in cytotoxic T lymphocytes (Davis et al., 2012). Other lysosomal Ca^2+^ channels, such as TRPML1, may also control cytotoxic T lymphocyte degranulation, as suggested in NK cells (Goodridge et al., 2019). Our Ca^2+^ measurements show that chronic acidosis enlarges both ER- and lysosome-releasable Ca^2+^ pools, with no effect on basal cytosolic Ca^2+^. A recent study in cell line reported that a subset of extracellular vesicles (EVs) carries LAMP1 at their surface, characterizing them as lysosome-derived EVs, which may include lysosomal cargo (Shen et al., 2026). The convergence of enlarged Ca^2+^ stores and expanded fusion machinery may synergistically lower the threshold for spontaneous lysosome-derived exosomes secretion and exocytosis. Importantly, the untargeted exocytosis was selective for the lysosome-derived organelles, as LAMP1 was increased in acidic condition while exosome markers CD9 and CD81 decreased, ruling out nonspecific vesicular shedding. This selectivity mirrors what is known about Ca^2+^ triggered SNARE-dependent fusion of secretory lysosomes (Hessvik and Llorente, 2018).

In essence, chronic acid adaptation expands and stabilizes the lysosomal membrane compartment while depleting its lytic cargo, uncoupling lysosomal abundance from cytotoxic capacity. The downstream consequences of this lysosomal remodeling are twofold. First, the accumulation of lysosomal membrane proteins, including LAMP1, LAMP2, CD63, CD68, without corresponding increase in their transcripts, suggests post-transcriptional stabilization, possibly through reduced membrane turnover or altered endolysosomal trafficking. In cancer cells, chronic acid exposure results in increased levels of LAMP2 to protect lysosomal membranes from acid proteolysis (Damaghi et al., 2015), and our data in CTLs align with this observation. Notably, surface LAMP1 also protects cytotoxic lymphocytes from degranulation-associated self-damage (Cohnen et al., 2013), whether the cathepsin-dependent loss of surface LAMP1 in acid-cultured CTLs (**Figure 4G**) compromises this protection remains to be tested. Second, the hyperproteolytic lysosomal environment degrades intraluminal cargo, depleting granzyme B from secretory lysosomes. In cytotoxic lymphocytes, the best-characterized roles of lysosomal proteases are activating rather than degradative: cathepsins C and H process pro-granzymes into their mature forms (D’Angelo et al., 2010; Pham and Ley, 1999), and cathepsin L contributes to perforin maturation (Konjar et al., 2010). A previously proposed cytoprotective role for surface cathepsin B during degranulation (Balaji et al., 2002) was not supported by genetic evidence, as cathepsin B–deficient CTLs survive target-cell killing normally (Baran et al., 2006). A degradative action of cathepsins toward lytic cargo, as proposed here, has not been described. To our knowledge, we provide the first evidence for a cathepsin B switch in cytotoxic lymphocytes, resulting in enhanced cargo degradation within a compartment that simultaneously undergoes constitutive, untargeted exocytosis, thus impairing the tumor killing capacity of CTLs. Consistent with this model, pharmacological inhibition of cysteine cathepsins increased intracellular granzyme B and increase surface LAMP1 exposure in CTLs (**Figure 4F, G**), implicating possible role of cathepsin B and, or L activity in both the cargo and membrane phenotypes of acid adaptation. Whether the reduced activation-induced surface pool and the constitutive shedding of LAMP1-positive vesicles impair the survival of acidic CTLs after degranulation remains to be tested. Beyond these membrane and cargo phenotypes, the lysosomal remodeling we observe, including luminal hyper-acidification, may also impinge on metabolic control, as the lysosome serves as a metabolic signaling hub, indeed, mTORC1 activation occurs at the lysosomal surface(Zoncu et al., 2011), is suppressed in acidic CD8⁺ T cells, together with reduced c-Myc levels (Vuillefroy de Silly et al., 2024). Yet, our proteomics revealed an enrichment of oxidative phosphorylation–related proteins in chronically acid-cultured CTLs and a positive trend toward an increase of MYC levels upon re-stimulation, suggesting that chronic acidosis modulates MYC activity rather than simply reducing MYC protein abundance.

These findings have translational implications for adoptive cell therapy. CAR-T cells targeting solid tumors face the same sustained acidosis, and the lytic defect begins before target encounter as the acid environment continuously drains lytic granules. Our data suggest two complementary therapeutic strategies. First, modulation of lysosomal hydrolysis: in the present study, cysteine cathepsin inhibition preserved intracellular granzyme B and restored surface LAMP1 in acid-cultured CTLs (**Figure 4F, G**), and V-ATPase modulators could act further upstream by limiting luminal hyperacidification. Whether cathepsin inhibition also restores target-cell killing under acidic conditions remains to be tested. Second, systemic pH buffering with oral bicarbonate improves anti-tumor immunity and synergizes with checkpoint blockade in preclinical models (Pilon-Thomas et al., 2016; Robey et al., 2009), and our work provides a mechanistic basis for this effect at the organelle level: acidic TME neutralization would halt constitutive exocytosis by restoring normal lysosomal Ca^2+^ homeostasis. A caveat is that our chronic acidification model uses PIPES-buffered medium, which does not recapitulate the full complexity of the tumor interstitial fluid. However, pH neutralization alone restores CTL function even in the presence of lactate (Mendler et al., 2012), supporting the physiological relevance of an acid-only model. We have not yet tested whether the constitutive exocytosis and cargo depletion occur in vivo in tumor-infiltrating CTLs, although the LAMP1-high/granzyme B-low phenotype observed here is reminiscent of dysfunctional CD8⁺ T cell states in human tumors (Thommen and Schumacher, 2018; Zheng et al., 2021).

Together, these data identify the lysosome as a principal site of adaptation in human CD8⁺ T cells exposed to chronic extracellular acidity. Acid-cultured CTLs hyperacidify their lysosomal lumen and remodel its protein composition, accumulating lysosomal membrane proteins and hydrolase activity while losing granzyme B. In parallel, chronic acidity enlarges the releasable ER and lysosomal Ca^2+^ stores without altering basal cytosolic Ca^2+^. Acid-exposed CTLs release lysosomal cargo constitutively, in the absence of target engagement, and inhibiting cathepsin B preserves their granzyme B stores. These observations link acid-driven lysosomal remodeling to a loss of cytotoxic capacity that begins before the target is encountered and point to lysosomal proteolysis as the step at which it can be interrupted.

## Materials

### Cell culture and acidic medium preparation

The human acute lymphoblastic leukemia NALM6-GFP cell line was cultured in RPMI with Glutamax (Gibco, #61870036) supplemented with 10% fetal bovine serum (FBS) and 1% penicillin/streptomycin (P/S). The HEK293-PG13-CD19-CAR retroviral packaging cell line was maintained in DMEM with Glutamax (Gibco #31966-021) supplemented with 10% FBS and 1% P/S (named complete DMEM).

Human CTLs were cultured in RPMI with Glutamax supplemented with 10% FBS 1% P/S, 100 µM non-essential amino acids (Gibco #11140-050), 1 mM sodium pyruvate (Gibco #11360-039) (named complete RPMI) and 20 ng/ml of IL-2. IL-2 was produced as previously described (Gaggero et al., 2022).

Acidic medium was prepared by supplementing complete RPMI with IL-2 (20 ng/mL) 12 mM HCl 37% (VWR #20255.290) and 20 mM PIPES pH 6.5 (ThermoFisher #J61224.AK) maintaining the medium at pH 6.5 for at least 3 days.

### Isolation and activation of human CTLs

Human CD8^+^ lymphocytes were purified from peripheral blood mononuclear cells (PBMCs) from buffy coats of healthy donors (Etablissement Français du Sang). PBMCs were isolated by Ficoll density gradient, then 2 × 10^8^ PBMCs were incubated 15 min at 4°C with 15 µl of anti-CD8 FITC antibody (Biolegend, #300906). After washing with MACS buffer, cells were incubated 15 min at 4°C with 70 µl of anti-FITC beads (Miltenyi #130-048-701). Finally, human CD8^+^ positive cells were isolated by magnetic separation using an LS column (Miltenyi #130-042-401).

Cells were plated at 2×10^6^ cell/ml and activated in the presence of T cell Transact (Miltenyi #130-111-160) using 10 µl per 10^6^ cells for 48 h.

### Retroviral production

5 × 10^6^ HEK293-PG13-CD19-CAR cells were plated in a T175 flask in 20 ml of complete DMEM. The day after the medium was replaced with 20 ml of complete DMEM without P/S. 24 h later, 10 ml of supernatant was collected and replaced with complete DMEM media without P/S. The supernatant was centrifuged (1 500 rpm, 5 min) to remove cell debris and stored at −80°C. The day after, the same steps were repeated with the remaining cell supernatant.

### Transduction of human CTL cells

1×10^6^ of human CTLs were centrifuged and resuspended in 1 ml of virus supernatant supplemented with 4 µg of polybrene (Merck, #TR-1003-G). Resuspended cells were spinoculated in a 24 well plate for 2 h at 2000g at 32°C. After the spinoculation, cells were incubated 6 h at 37°C 5% CO2, then 800 µl of the supernatant were replaced with complete RPMI containing 20 ng/ml of IL-2. Three and seven days later, transduction efficiency was monitored by flow cytometry.

### Long term culture of human CD8+ T cells in acidic conditions

After activation or transduction, human CTLs or CAR-T cells were incubated in 12-well plates and maintained at 0.5-1×10^6^ cell/ml either in physiological (pH 7.5) or acidic (pH 6.5) complete RPMI supplemented with IL-2 (20 ng/ml). The medium was refreshed every 3 days to maintain acidic conditions. After 7 days of culture, cells were harvested and analyzed by flow cytometry or used for further functional assays. CTLs cultured at pH 7.5 and 6.5 are referred as neutral and acidic CTLs respectively.

### Sample staining for flow cytometry analysis

To assess viability, cells were incubated with the viability dye Zombie NIR (BioLegend #423105) diluted at 1:1000 in PBS for 20 minutes at 4°C, and the reaction was stopped by adding MACS buffer. Next, cells were incubated 15 minutes at RT with a mix of antibodies diluted at 1:100 in MACS buffer for surface markers, then washed twice with MACS buffer.

Transduction efficiency was evaluated with Withlow/218 Linker antibody (Clone E3U7Q, CST, 1:200, 72160S).

For cytokine production assay, the viability staining was performed as described above. Thereafter, cells were fixed and permeabilized by using the BD Cytofix/Cytoperm kit (BD Biosciences #554714). Then, cells were incubated with anti-human TNF PE-Dazzle (BioLegend #502946, 1/1000), anti-human IFNg APC (Biolegend, #506510, 1/1000), anti-human IL-2 BV410 (BioLegend #500328, 1/1000), and anti-human Granzyme B (BD Bioscience, #560213, 1/1000) overnight at 4°C in Perm/Wash buffer included in the kit.

For LAMP1 intracellular staining, viability staining was performed as described above. Cells were fixed and permeabilized by using the BD Cytofix/Cytoperm kit (BD Biosciences #554714). Then, cells were incubated with anti-human LAMP1/CD107a PE (BioLegend #328608, 1/100) 30 min at 4°C in Perm/Wash buffer included in the kit.

All flow cytometry was performed on a Cytoflex XL (Beckman), and analyses were performed using FlowJo software (version 10.10).

### Cytotoxic assay

Flat-bottom 96 well plates were coated for 1h at RT with 50 µl of bovine fibronectin (Sigma, F1141-5MG) at 5 µg/ml. After the incubation, the solution was removed and wells were dried for 30 min. NALM6-GFP cells and control CTLs or CAR-T cells (neutral and acidic counterparts) were mixed at an effector/target ratio of 1:1 in the different pH media. The co-culture solution was added to the wells and the plate was incubated in the IncuCyte SX5 live-cell analysis system (Sartorius) for 3 days with an acquisition every 2h. Target cell lysis was reflected by the decrease in GFP signal normalized to the initial signal at the beginning of the coculture.

### Cytokine production assays

100 000 CTLs were incubated in 96 well plates at physiological or acidic pH and were stimulated with Cell Stimulation Cocktail with protein transport inhibitors (eBioscience, 00-4975-03) for 4 h. After stimulation, cells were washed and stained for viability and selected markers.

### Degranulation assay

In 96 well plates, 100,000 CTLs were stimulated in the different pH conditions with Cell Stimulation Cocktail with protein transport inhibitors (eBioscience, 00-4975-03) for 4 h in the presence of anti-LAMP1/CD107a antibody (PE, 1:100, BioLegend #328608). After stimulation, cells were stained for viability and selected markers.

Neutral or acidic CAR-T cells were co-cultured with NALM6-GFP+ cells at an effector:target ratio of 1:2 in the presence of anti-LAMP-1/CD107a antibody (PE, 1:100, BioLegend #328608). The medium was supplemented with 1/1000 BrefeldinA (BD Bioscience, #51-2301KZ) and Monensin (BD Bioscience, #51-2092KZ). After 4 h, cells were harvested and stained for viability and selected markers.

### Lysotracker staining

Neutral and acidic CTLs were incubated for 2h at physiological pH with 70 nM LysoTracker Deep Red (Thermo Fisher, #L12492). After incubation, cells were washed and analyzed by flow cytometry.

### LysoSensor staining

Human CD8^+^ T cells were isolated by magnetic separation using LS column (Miltenyi #130-042-401). Cells were plated at 2×10^6^ cell/ml and activated in the presence of T cell Transact (Miltenyi #130-111-160) using 10 µl per 10^6^ cells for 48 h. After activation, human CD8^+^ T cells were incubated in 12-well plates and maintained at 0.5-1×10^6^ cell/ml either in physiological (pH 7.5) or acidic (pH 6.5) complete RPMI supplemented with IL-2 (20 ng/ml). The medium was refreshed every 3 days to maintain acidic conditions. After 3 days of culture, cells were attached to coverslips by incubating them at 37°C for 1 hour in PBS, then PBS was replaced with complete RPMI at neutral pH containing LysoSensor Yellow/Blue DND160 (ThermoFisher, L7545) or Lysosensor Yellow/Blue DND160 (ThermoFisher, L7545) following the manufacturer’s instructions. Images were acquired using a LSM700 confocal microscope (Leica) or a Zeiss Apotome 3. Quantitative analyses of the images were performed using ImageJ software.

### Ca^2+^ flux assay

Neutral and acidic CTLs were loaded with the ratiometric dye indo-1 AM (Thermo Fisher Scientific, Waltham, MA, USA) as a Ca²⁺ indicator. 700 000 cells per milliliter were incubated with 0.7 μM indo-1 AM in the corresponding medium for 30 min at 37 °C in the dark and were then washed. The evolution of intracellular calcium concentration ([Ca²⁺]i) was measured every second as the fluorescence ratio F400/F475 (RF400/F475) using a UV-equipped flow cytometer. Baseline [Ca²⁺]i was recorded in a 0 mM Ca²⁺ solution containing 140 mM NaCl, 5 mM KCl, 1 mM MgCl₂, 5 mM glucose, and 10 mM HEPES (pH 7.4). After 120s, cells were treated with either 1 μM thapsigargin, a sarco/endoplasmic reticulum Ca²⁺-ATPase (SERCA) inhibitor (Focus Biomolecules), 50 μM GPN (Thermo Fisher, #J64718.MC) or 10 µM ionomycin (Thermo Fisher Scientific, Waltham, MA, USA). After 300 s of treatment with thapsigargin or GPN, ionomycin was added to the cells. Fluorescence was monitored using an LSRFortessa X-20 flow cytometer, and median fluorescence values (n = 500-1000 cells per time point) were extracted for further analysis. Calcium storage was calculated as previously described [Lewuillon et al., 2022].

### Cathepsin protease activity assay

Whole cell lysates were prepared in 200mM Na^+^ citrate buffer pH 4.0 containing 1% Triton X-100. Cysteine lysosomal protease activities were measured using 10-20 μg of total protein in 200 μL of assay buffer (200mM Na^+^ citrate buffer) containing 10 μM Omnicathepsin substrate [Z-Phe-Arg-AMC] (Enzo Life Sciences #BML-P139) or 10 μL cysteine protease substrate [Z-Arg-Arg-AMC]. AMC release was quantified at 460 nm in a fluorescent plate reader (Varioskan Lux, ThermoFisher). Activities were calculated as arbitrary units.

### Cysteine cathepsin inhibition

CTLs were incubated for 3 days at pH 7.5 or pH 6.5 in the presence or absence of the cell-permeant cysteine cathepsin inhibitor Ca074-Me (10 µM; Selleck Chemicals, #S7420), which inactivates cathepsins B and L in intact cells (Buttle et al., 1992; Montaser et al., 2002). Cells were then harvested, stained with Zombie NIR viability dye, and stained for intracellular granzyme B as described above. In parallel, surface LAMP1 was measured after stimulation by Cell Stimulation Cocktail with protein transport inhibitors (eBioscience, 00-4975-03) on live cells by flow cytometry as described above.

### Western Blot

Whole cell extracts were prepared using RIPA lysis buffer (50 mM Tris-HCl pH 7.8, 150 mM NaCl, 0.5% sodium deoxycholate, 0.1% SDS, 1% NP-40) supplemented with a 1:100 dilution of protease inhibitor cocktail (Roche, #11836170001) and phenylmethylsulfonyl fluoride (ITW Reagents, #A0999). Equal amounts of protein were denatured and separated by SDS-PAGE on 12% or 15% polyacrylamide gels, then transferred to 0.20-µm nitrocellulose membranes (Bio-Rad, #10484059). Membranes were blocked with 5% non-fat dry milk in PBST (1X PBS, 0.1% Tween-20) for 1 hour at RT. Membranes were subsequently incubated overnight at 4 °C with primary antibodies diluted 1:1,000 in blocking buffer. Following three 5-minute washes with PBST, membranes were incubated for 1 hour at RT with the appropriate horseradish peroxidase (HRP)-conjugated secondary antibodies (1:5,000). Protein bands were visualized using Clarity Western ECL Substrate (Bio-Rad, #170-5061) and imaged on a ChemiDoc Imaging System (Bio-Rad). Primary antibodies used were: anti-CD81 (Invitrogen, #10630D), anti-CD9 (Invitrogen, #10626D), anti-CtsL (R&D Systems, #MAB9521), anti-CtsB (Abcam, #ab9295), anti-CtsD (Cell Signaling Technology, #2284), anti-LAMP1 (Abcam, #ab24170), anti-β-actin (Santa Cruz Biotechnology, #sc-47778), and anti-LAMP2 (Abcam, #ab25631), anti-TFEB (Bethyl, #A303-673A), Histone 3 (Abcam, #ab1791). HRP-conjugated secondary antibodies were: anti-mouse IgG (Cell Signaling Technology, #7076), anti-rabbit IgG (Cell Signaling Technology, #7074), anti-rat IgG (Cell Signaling Technology, #7077S), and anti-sheep IgG (Jackson ImmunoResearch, #713-035-003).

### Subcellular fractionation

Subcellular proteins were extracted using the ProteoExtract Subcellular Proteome Extraction Kit (#539790, Sigma-Aldrich) according to the manufacturer’s instructions. Cultured cells were harvested by trypsinization (5-10 min), pelleted (300 g, 10 min), washed twice in 800 µL ice-cold Wash Buffer with a 5-min incubation at 4 °C under gentle rotation to prevent clumping, and re-pelleted (300 g, 10 min, 4 °C). For the cytosolic fraction (Fraction 1), the pellet was resuspended in 300 µL ice-cold Extraction Buffer I supplemented with 1.5 µL Protease Inhibitor Cocktail (PIC; #P8340, Sigma-Aldrich), incubated for 10 min at 4 °C with gentle agitation, and centrifuged (1000 g, 10 min, 4 °C). For the membrane/organelle fraction (Fraction 2), the remaining insoluble pellet was incubated in 300 µL ice-cold Extraction Buffer II with 1.5 µL PIC for 30 min at 4 °C and centrifuged (6000 g, 10 min, 4 °C). For the nuclear fraction (Fraction 3), the pellet was resuspended by gentle pipetting in 150 µL Extraction Buffer III containing 1.5 µL PIC and 0.45 µL Benzonase nuclease, incubated for 10 min at 4 °C, and centrifuged (6800 g, 10 min, 4 °C). All supernatants were collected separately, stored on ice or at −80 °C, and subsequently processed for Western blot analysis.

### Proteomic sample preparation analysis

For basal samples, neutral and acidic CTLs were harvested after 7 days of culture on their respective pH medium.

For reactivated samples, neutral and acidic CTLs were reactivated with T cell Transact (Miltenyi #130-111-160, 1/500) for 4 h either at pH 7.5 or pH 6.5. Cells were washed in PBS and pelleted at 5,000g for 1 min at 4°C.

Protein concentrations were determined using a micro-BCA assay from lysates prepared in 5% SDS and 50 mM TEAB. 100 µg of protein were processed using S-Trap Micro spin columns, according to the manufacturer’s instructions with minor modifications. Disulfide bonds were reduced with 10 mM DTT for 10 min at 95°C, followed by alkylation with 40 mM iodoacetamide (IAA) for 30 min in the dark. Samples were then loaded onto S-Trap spin columns. Trapped proteins were washed five times with S-Trap binding buffer using 500 µL per wash.

Proteins were digested with trypsin (Pierce, Thermo Fisher Scientific) at an enzyme-to-protein ratio of 1:40 using a two-step digestion protocol. Samples were digested overnight at 37 °C in 160 µL of 50 mM TEAB, followed by a second digestion for an additional 6 h at 37 °C. Peptides were eluted with 160 µL of 50 mM TEAB, followed by 160 µL of 0.2% aqueous formic acid and 160 µL of 50% acetonitrile containing 0.2% formic acid, with centrifugation at 1,000 × g for 1 min after each elution step. Tryptic peptides were dried in a SpeedVac and stored at −20 °C before LC-MS analysis.

### LC-MS Analysis

Peptides (equivalent of 200 ng) were injected onto a nanoscale C18 reverse-phase chromatography system (Vanquish, Thermo Scientific) and electrosprayed into an Astral Orbitrap Mass Spectrometer (Thermo Fisher). For liquid chromatography the following buffers were used: buffer A and weak buffer wash (0.1% formic acid in Milli-Q water (v/v) buffer B and strong buffer wash (80% acetonitrile and 0.1% formic acid in Milli-Q water (v/v). Samples were loaded onto a trap column (Pep Map Neo C18, 5µm, 300µm x5 mm) Thermo Scientific) equilibrated in buffer A with the following loading settings, mode: flow control, flow 10µl/min and loading volume set to automatic. The wash and equilibration settings were as follows equilibration factor 2, trap column, fast wash and equilibration on, zebra wash on, zebra wash cycle 4, equilibration factor automatic, mode combined control, flow 10µl/min, pressure 800 bar. The column (Pep MAP C18, 2µm, 150 µm x15 cm) was equilibrated with 4% buffer and peptides were eluted at a variable flow rate, starting at 1.3 µl/min to 0.8 µl/min from 4% buffer B to 8% buffer B in 0.1 min, then from 8% buffer B to 22.5% buffer B in 13 min, and from 22.5% b to 35% B in 6.90 min. The low rate is increased from 0.8 µl/min to 2 µl/min in 0.4 min with buffer B increasing from 35% to 55%. The column is finally washed at a flow of 2µl/min with 99% buffer B. The column was kept at a constant temperature of 50°C.

The data was acquired using an easy spray source operated in positive mode with spray voltage at 2.0 kV, and the ion transfer tube temperature at 280°C. The MS was operated in DIA mode. A scan cycle comprised a full MS scan (m/z range from 380-980), with orbitrap resolution at 2400000, RF lens at 40%, AGC target set to custom, normalised AGC target at 500%, absolute AGC value set to 5.00e6, maximum injection time 5ms and micros-can set to 1. MS scan data was acquired in profile mode.

MS survey scan was followed by MS/MS DIA scan events using the following parameters: DIA window type set to auto, isolation window 4, window overlap set to 0, window placement optimisation on, number of scan events 150, collision energy type normalized, HCD collision energy 25%, detector type Astral, scan range 150-2000, normalized AGC target 500%, absolute AGC target 5.000e4, maximum injection time 3 ms, microscan set to 1, loop control time, time 0.6s. The DIA data was collected in centroid mode.

### Proteomic data analysis

Raw mass spectrometry data was searched with Spectronaut (v19.5.241126.62635, Biognosys) with a human combined SwissProt/TremBL FASTA and a contaminants FASTA file using a library free method. Peptide and protein false discovery rate was set to 0.01, and calculated with an inverse decoy library. Minimum peptide length was set to 7 and the number of missed cleavages was set to 2, with carbaminodylation set as a fixed modification and N-terminal acetylation and methionine oxidation set as variable modifications. Quantification was set to QUANT 2.0 and cross-run normalisation was switched off.

Protein copy numbers were calculated using the proteomic ruler in Perseus, p values, and log2 fold changes were calculated using RStudio (v.2026.08.01+195) with the Bioconductor package ‘limpa’. Proteins were considered differentially abundant at unadjusted p value < 0.05 and absolute fold-change > 1.5. Pathway enrichment was done using ‘clusterProfiler’.

Concordance between RNA and protein regulation was assessed by comparing genes meeting the above thresholds in each dataset.

A TFEB target gene signature (n = 1,425 genes; MSigDB C3:TFT:GTRD collection,(Yevshin et al., 2019)) was analyzed by Gene Set Enrichment Analysis using the fgsea package (v.1.28.0).

### RNA-Seq analysis

Neutral and acidic CTLs were pelleted and RNA was extracted using a Quick-RNA Microprep kit (Zymo Research, #R1051) the following manufacturer’s instructions. Sample processing and quantification were performed by Plasmidsaurus. Quality of the fastq files was assessed using FastQC v0.12.1. Reads were then quality filtered using fastp v0.24.0 with poly-X tail trimming, 3’ quality-based tail trimming, a minimum Phred quality score of 15, and a minimum length requirement of 50 bp. Quality-filtered reads were aligned to the reference genome using STAR aligner v2.7.11 with non-canonical splice junction removal and output of unmapped reads, followed by coordinate sorting using samtools v1.22.1. PCR and optical duplicates were removed using UMI-based deduplication with UMIcollapse v1.1.0. Alignment quality metrics, strand specificity, and read distribution across genomic features were assessed using RSeQC v5.0.4 and Qualimap v2.3, with results aggregated into a comprehensive quality control report using MultiQC v1.32. Gene-level expression quantification was performed using featureCounts (subread package v2.1.1) with strand-specific counting, multi-mapping read fractional assignment, exons and three prime UTR as the feature identifiers, and grouped by gene_id. Final gene counts were annotated with gene biotype and other metadata extracted from the reference GTF file. Differential expression analysis was performed using DESeq2.

### Statistical analysis

Data are presented as means ± SEM of at least three independent experiments, and paired two-tailed t-tests were performed unless otherwise stated after confirming normal distribution of samples with the Shapiro-Wilk test. Multiple group comparisons were performed using one-way analysis of variance (ANOVA) unless otherwise stated. ns, not significant; *p < 0.05, **p < 0.01, ***p < 0.001, and ****p < 0.0001. All analyses were performed using Prism 9 software (GraphPad).

## Acknowledgments

The authors acknowledge Prof Doreen Cantrell (University of Dundee) for her help in designing and interpret proteomic analysis, Dr Claire Hivroz (Insitut Curie) for her help in reviewing the manuscript. The authors thank Nathalie Jouy and Emilie Floquet from Plateformes Lilloises en Biologie et Santé (PLBS) - UAR 2014 - US 41 hosting the flow core facility. We thank FingerPrints Proteomics Facility, University of Dundee for proteomic sample preparation and mass spectrometry analysis.

We thank the ONCOLille institute.

## Contribution

Conceptualization: BS, JM-F, SM. Methodology: BS, RC, CR, ST, YT, LS, JM-F, SM. Investigation and data analysis BS, RC, CR, ST, AC, AK, EP, LM, SG, RR, RE, I.M and BQ. Supervision: JM-F, SM. Funding acquisition: JM-F, SM. Writing – original draft: SM and BS, JM-F. Review & editing: all authors.

